# Optimal time frequency analysis for biological data - pyBOAT

**DOI:** 10.1101/2020.04.29.067744

**Authors:** Gregor Mönke, Frieda A. Sorgenfrei, Christoph Schmal, Adrián E. Granada

## Abstract

Methods for the quantification of rhythmic biological signals have been essential for the discovery of function and design of biological oscillators. Advances in live measurements have allowed recordings of unprecedented resolution revealing a new world of complex heterogeneous oscillations with multiple noisy non-stationary features. However, our understanding of the underlying mechanisms regulating these oscillations has been lagging behind, partially due to the lack of simple tools to reliably quantify these complex non-stationary features. With this challenge in mind, we have developed pyBOAT, a Python-based fully automatic stand-alone software that integrates multiple steps of non-stationary oscillatory time series analysis into an easy-to-use graphical user interface. pyBOAT implements continuous wavelet analysis which is specifically designed to reveal time-dependent features. In this work we illustrate the advantages of our tool by analyzing complex non-stationary time-series profiles. Our approach integrates data-visualization, optimized sinc-filter detrending, amplitude envelope removal and a subsequent continuous-wavelet based time-frequency analysis. Finally, using analytical considerations and numerical simulations we discuss unexpected pitfalls in commonly used smoothing and detrending operations.

## 1 Introduction

Oscillatory dynamics are ubiquitous in biological systems. From transcriptional to behavioral level these oscillations can range from milliseconds in case of neuronal firing patterns, up to years for the seasonal growth of trees or migration of birds (Goldbeter et al. [2012], Gwinner [2003], Rohde and Bhalerao [2007]). To gain biological insight from these rhythms, it is often necessary to implement time-series analysis methods to detect and accurately measure key features of the oscillatory signal. Computational methods that enable analysis of periods, amplitudes and phases of rhythmic time series data have been essential to unravel function and design principles of biological clocks (Lauschke et al. [2013], Ono et al. [2017], Soroldoni et al. [2014]). Here we present `pyBOAT`, a framework and software package with a focus on usability and generality of such analysis.

Many time series analysis methods readily available for the practitioner rely on the assumption of stationary oscillatory features, i.e. that oscillation properties such as the period remain stable over time. A plethora of methods based on the assumption of stationarity have been proposed which can be divided into those working in the frequency domain such as Fast Fourier transforms (FFT) or Lomb-Scargle periodograms (Lomb [1976], Ruf [1999]) and those working in the time domain such as autocorrelations (Westermark et al. [2009]), peak picking (Abraham et al. [2018]) or harmonic regressions (Edwards et al. [2010], Halberg et al. [1967], Naitoh et al. [1985], Straume et al. [1991]). In low noise systems with robust and stable oscillations, these stationary methods suffice to reliably characterize oscillatory signals. Recordings of biological oscillations frequently exhibit noisy and time-dependent features such as drifting period, fluctuating amplitude and trend. Animal vocalization (Fitch et al. [2002]), temporal changes in the activatory pathways of somitogenesis (Tsiairis and Aulehla [2016]) or reversible and irreversible labilities of properties in the circadian system due to aging or environmental factors (Pittendrigh and Daan [1974], Scheer et al. [2007]) are typical examples where systematic, often non-linear changes in oscillation periods occur. In such cases, the assumption of stationarity is unclear and often not valid, thus the need to use nonstationary-based methods that capture time-dependent oscillatory features. Recently, the biological data analysis community has developed tools that implement powerful methods tailored to specific steps of time-series analysis such as rhythmicity detection (Hughes et al. [2010], Thaben and Westermark [2014]), de-noising and detrending, and the characterization of nonstationary oscillatory components (Leise [2013], Price et al. [2008]).

To extract time-dependent features of non-stationary oscillatory signals, methods can be broadly divided into those that rely on operations using a moving time-window (e.g. wavelet transform) and those that embeds the whole time series into a phase space representation (e.g. Hilbert transform). These two families are complementary, having application-specific advantages and disadvantages, and in many cases both are able to provide equivalent information about the signal (Quiroga et al. [2002]). Due to the inherent robustness in handling noisy oscillatory data and its interpretability advantages, we implemented at the core of `pyBOAT` a continuous-wavelet-transform approach.

As a software package `pyBOAT` combines multiple steps in the analysis of oscillatory time series in an easy-to-use graphical user interface that requires no prior programming knowledge. With only two user-defined parameters, `pyBOAT` is able to proceed without further intervention with optimized detrending, amplitude envelope removal, spectral analysis, detection of main oscillatory components (ridge detection), oscillatory parameters readout and visualization plots (Figure 1A). `pyBOAT` is developed under an open-source license, is freely available for download and can be installed on multiple operatings systems.

**Figure 1:**
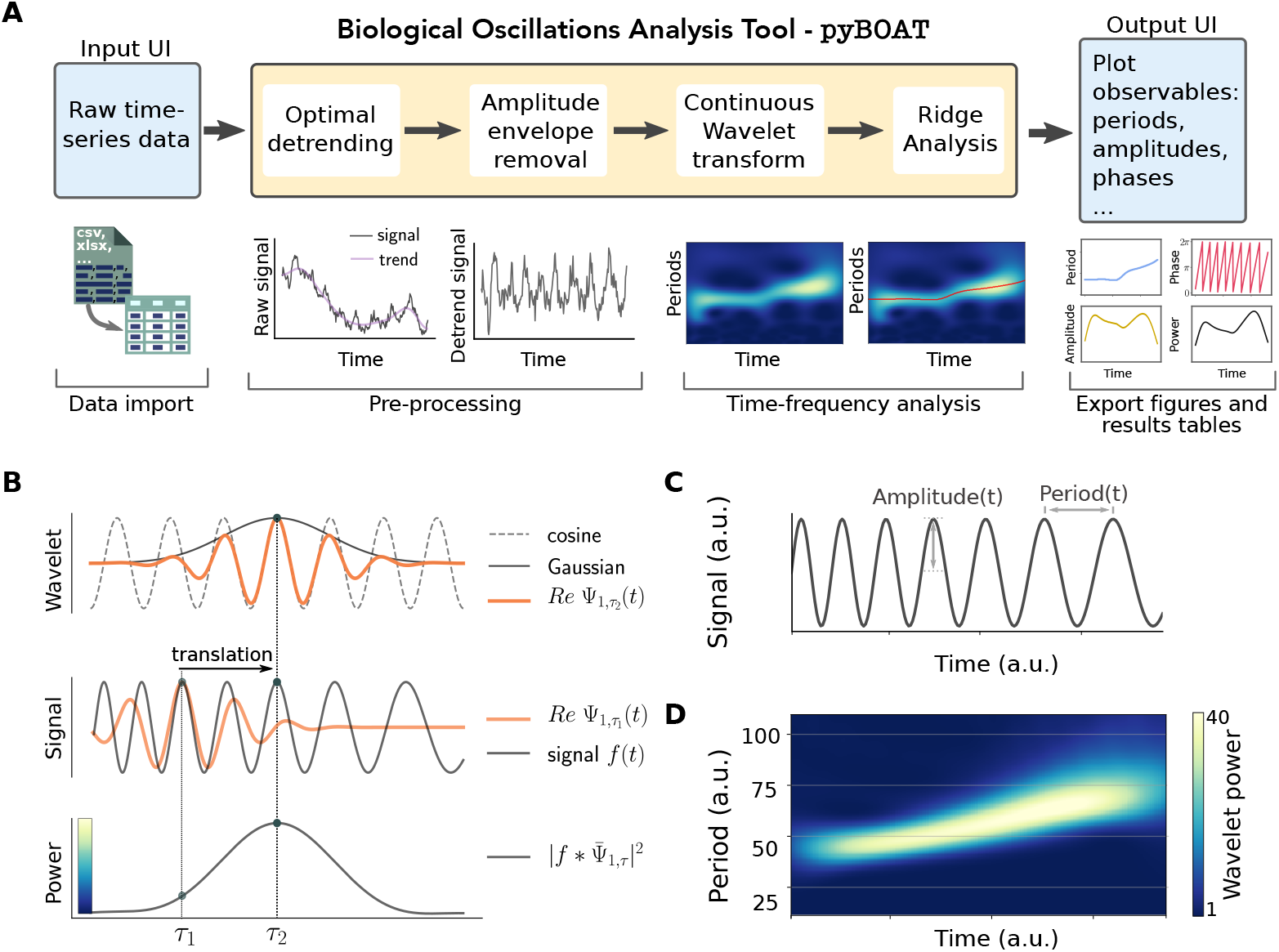
Overview of the workflow implemented in pyBOAT. A) Schematic showing the proposed and implemented analysis steps. B) *Top panel*: A cosine function (dotted gray line) multiplied by a Gaussian envelope (bold black line) defines the Morlet Wavelet (orange line), here depicted for scale *s* = 1 and localized at *τ* = *t*_2_. *Middle panel*: The same Morlet Wavelet, translated in time with *τ* = *t*_1_, is shown together with a sweeping signal whose instantaneous period coincides with the Morlet of scale *s* = 1 exactly at *τ*_2_. *Bottom panel*: Result of the convolution with the sliding Morlet Ψ_1*,τ*_ (*t*) along signal *f*(*t*). The power quickly decreases away from *τ*_2_. The curve corresponds to one row in the Wavelet power spectrum of panel D). C) Synthetic signal with periods sweeping from *T*_1_ = 30s (*f*_1_ ≈ 0.033Hz) to *T*_2_ = 70s (*f*_2_ ≈ 0.014Hz). D) Wavelet power spectrum shows time-resolved (instantaneous) periods.

In the first section of this work we lay out the mathematical foundations at the core of `pyBOAT`. In the subsequent section 3 we describe artifacts generated by the widely used smoothing and detrending techniques and how they are resolved within `pyBOAT`. In section 4 we describe the theory behind spectral readouts in the special case of complex amplitude envelopes. We finalize this manuscript with a short description of the user interface and software capabilities.

## 2 Basic Wavelet Theory

In this section we aim to lay down the basic principles of wavelet analysis as employed in our signal analysis tool, albeit the more mathematical subtleties are moved to the Appendix.

### Introduction to the Wavelet transform

The classic approach to do frequency analysis of periodic signals is the well-known Fourier analysis. Its working principle is the decomposition of a signal *f*(*t*) into Sines and Cosines, known as basis functions. These harmonic components have no *localization in time* but are sharply *localized in frequency*: each harmonic component carries exactly one frequency effective everywhere in time. Thus, the straighforward Fourier analysis underperforms in cases of time-dependent oscillatory features, such as when the period of the oscillation changes in time (Figure 1C). The goal behind wavelets is to reach an optimal compromise between time and frequency localization (Gabor [1946]). Gabor introduced a Gaussian modulated harmonic component, also known as *Morlet wavelet*:

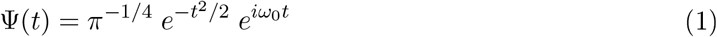

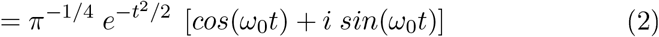

The basis harmonic functions for time-frequency analysis are then generated from the *mother wavelet* by *scaling* and *translation*:

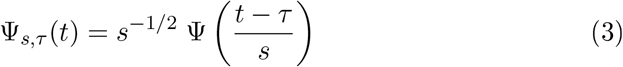

Varying the time localization *τ* slides the wavelet left and right on the time axis. Scale *s* changes the *center frequency* of the Morlet wavelet according to *ω_center_*(*s*) = *ω*_0_*/s* (see also Appendix equation (8)). Higher scales therefore generate wavelets with lower center frequency. The Gaussian envelope suppresses the harmonic component with frequency *ω_center_* farther away from *τ*, therewith localizing the wavelet in time (Figure 1B top panel). This frequency *ω_center_*(*s*) is conventionally taken as the Fourier equivalent (or pseudo-) frequency of a Morlet wavelet with scale *s*. It is noteworthy to state that wavelets are in general not as sharply localized in frequency as their harmonic counterparts (Figure S1). This is a trade-off imposed by the uncertainty principle to gain localization in time (Gröchenig [2013]).

The wavelet transform of a signal *f*(*t*) is given by the following integral expression:

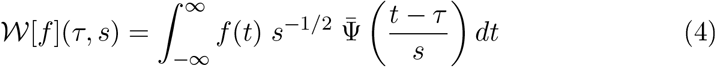

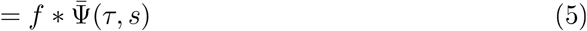

For a fixed scale, this equation has the form of a convolution as denoted by the ‘*’ operator, where 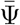 denotes the complex conjugate of Ψ. For an intuitive understanding it is helpful to consider above expression as the cross-correlation between the signal and the wavelet of scale *s* (or center frequency *ω_center_*(*s*)). The translation variable *τ* slides the wavelet along the signal. Since the wavelet decays fastly away from *τ*, only the instantaneous correlation of the wavelet with center frequency *ω_center_* and the signal around *τ* significantly contributes to the integral (Figure 1B middle and lower panel). By using an array of wavelets with different frequencies (or periods), this allows to scan for multiple frequencies in the signal in a time-resolved manner.

The result of the transform: 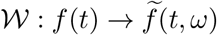 is a complex valued function of two variables, frequency *ω* and time localization *τ*. In the following, we implicitly convert scales to frequency via the corresponding central frequencies *ω_center_*(*s*) of the Morlet wavelets. To obtain a physically meaningful quantity, one defines the *wavelet power spectrum*:

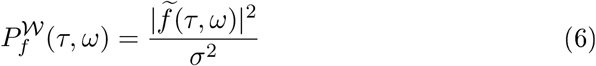

We adopted the normalization with the variance of the signal *σ*^2^ from Torrence and Compo [1998] as it allows for a natural and statistical interpretation of the wavelet power. By stacking the transformations in a frequency order, one constructs a two-dimensional time-frequency representation of the signal, where the power itself is usually color coded (Figure 1D), using a dense set of frequencies to scan for approximates of the *continuous Wavelet transform*.

### Wavelet Power Interpretation

It is important to note that the *time averaged* wavelet power spectrum

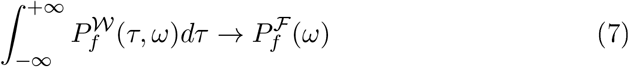

is an unbiased estimator for the true Fourier power spectrum 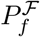 of a signal *f* (Percival [1995]).

This allows to directly compare Fourier power spectra to the wavelet power. White noise is the simplest noise process which may serve as a null hypothesis. Normalized by variance white noise has a flat mean Fourier power of one for all frequencies. Hence, also the variance normalized Wavelet power of one corresponds to the mean expected power for white noise: 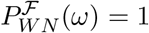 (Figure 2C). This serves as a universal unit to compare different empirical power spectra.

**Figure 2:**
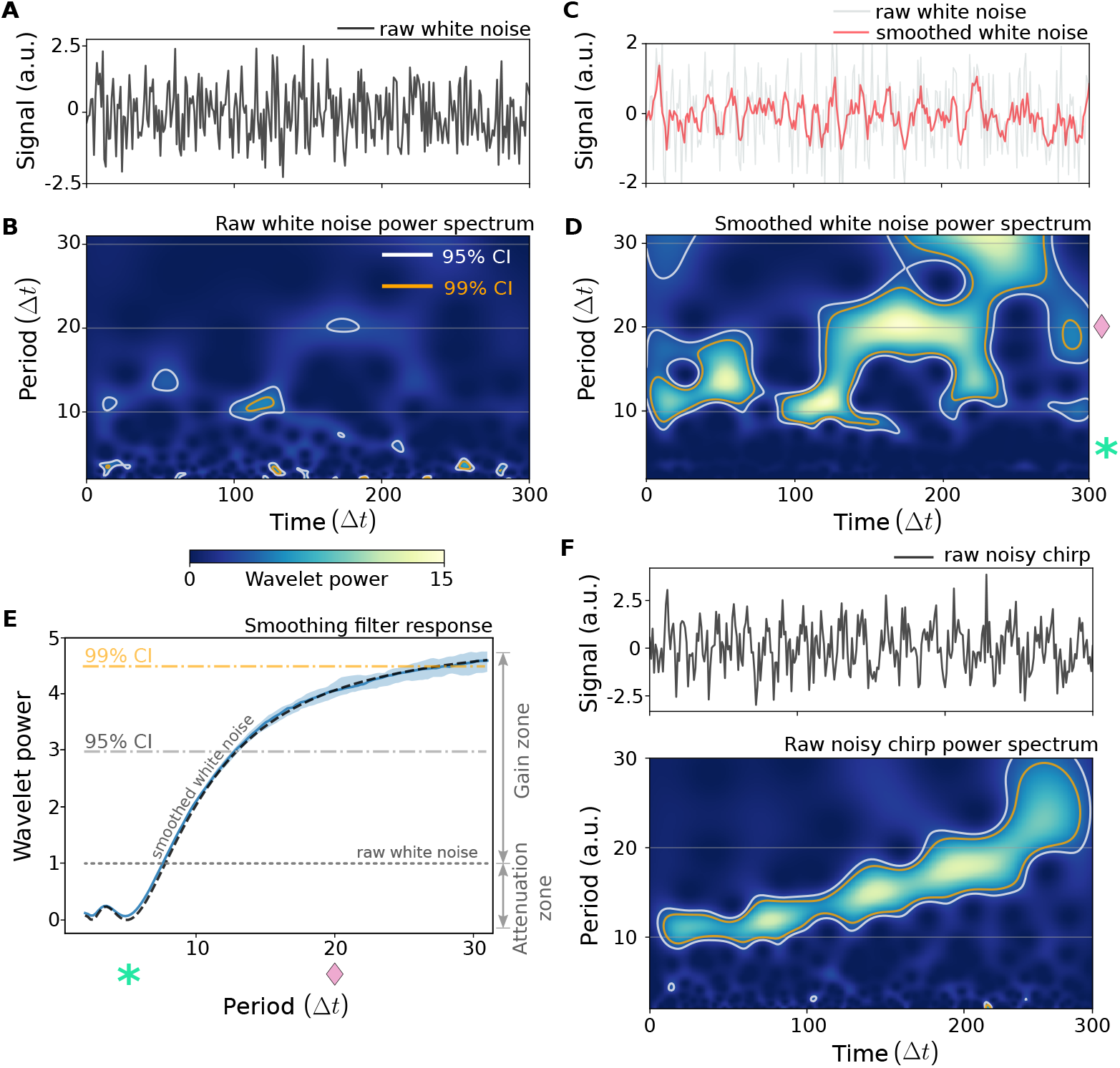
Smoothing inevitably introduces signal artefacts. A) Raw non-smoothed white noise time-series. B) Wavelet power spectrum of the raw white noise signal shown in A. 95%(white line) and and 99%(orange line) confidence contour levels are indicated. C) Moving average smoothed white noise signal (*red line*) from panel A. D) Wavelet power spectrum of the smoothed signal shown in C. Note the highly significant islands of high power 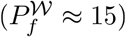 for periods greater than around 10 sampling intervals, the largest one is in the *T* ≈ 20Δ*t* band, marked by (♦). The (*) marks the end of the effective stop-band region of the filter. E) Attenuations and gain effects of smoothing with a moving average filter, (quartiles in blue). Low periods (high frequencies) are suppressed while higher periods are amplified up to 5-fold in average power. The *dashed black line* denotes the theoretically calculated Fourier spectrum of the moving average filter, scaled with 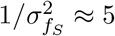, coinciding with the numerical results (blue shaded area). The gray dotted line depicts the mean power of a white noise signal. F) *Top:* Noisy slowing oscillation (chirp) time-series with a signal-to-noise ratio of 1. *Bottom:* Corresponding wavelet power spectrum, obtained from the raw unsmoothed signal as processed by `pyBOAT`. In panel E, the averaged wavelet power distribution was calculated using 50 time series with 25.000 sample points each and smoothed with a window size *M* = 5Δ*t*.

Extending these arguments to random fluctuations of the Fourier spectrum allows for the calculation of confidence intervals on wavelet spectra. If a background spectrum 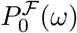 is available, the confidence power levels can be easily calculated as:

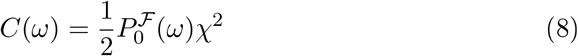

Assuming normality for the distribution of the complex Fourier components of the background spectrum, one can derive that the power itself is chi-square distributed (Chatfield [1995]). Thus, picking a desired confidence (e.g. *χ*^2^(95%) ≈ 6) gives the scaling factor for the background spectrum. Only Wavelet powers greater than this confidence level are then considered to indicate oscillations with the chosen confidence. The interested reader may find more details in section 4 of Torrence and Compo [1998].

A wavelet power of *C* = 3 corresponds to the 95% confidence interval in case of white noise (Figure 2B), which is frequency independent. For the practitioner, this should be considered the absolute minimum power, required to report ‘oscillations’ in a signal. It should be noted that especially for biological time series, due to correlations present also in non-oscillatory signals, white noise often is a poor choice for a null model (see also supplementary information A.3). A possible solution is to estimate the background spectrum from the data itself, this however is beyond the scope of this work.

## 3 Optimal Filtering - Do’s and Dont’s

A biological recording can be decomposed into components of interest and those elements which blur and challenge their analysis, most commonly noise and trends. Various techniques for *smoothing* and *detrending* have been developed to deal with these issues. Often overlooked is the fact that both, *smoothing* and *detrending* operations can introduce spectral biases, i.e. attenuation and amplification of certain frequencies. In this section we lay out a mathematical framework to understand and compare the effects of these two operations, showing examples of the potential pitfalls and at the same time providing a practical guide to avoid these issues. Finally, we discuss how `pyBOAT` minimizes most of these common artifacts.

### Smoothing

The operation which removes the fast, high-frequency (low period) components of a signal is colloquially called *smoothing*. This is most commonly done as a sliding time window operation (convolution). In general terms we can refer to a window function *w*(*t*) such that the smoothed signal is given as:

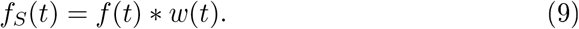

By employing the convolution theorem, it turns out that the Fourier transformation of the smoothed signal

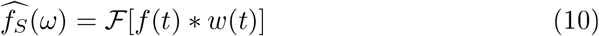

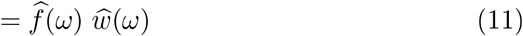

is simply given by the product of the individual Fourier transforms.

It follows that the Fourier power spectrum of the smoothed signal reads as:

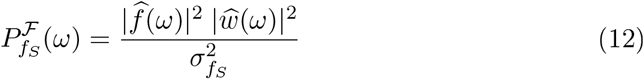

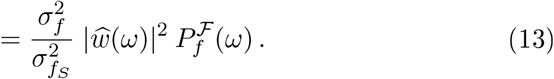

Applying a few steps of Fourier algebra shows that the original power spectrum 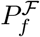 gets modified by the *low pass response* of the window function 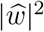 scaled by the ratio of variances 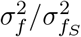. Also without resorting to mathematical formulas, smoothing and its effect on the time-frequency analysis can be easily grasped visually. A broad class of filtering methods falls into the category of *convolutional filtering*, meaning that there is some operation in a sliding window done to the data, e.g. for *moving average* and *LOESS* or *Savitzky-Golay* filtering (Savitzky and Golay [1964]). Moving average filter is a widely used smoothing technique, defined simply by a box-shaped window that slides in the time domain. In Figure 2 we summarize the spurious effects that this filter can have on noisy biological signals. White noise, commonly used as descriptor for fluctuations in biological systems, is a random signal with no dominant period. The lack of dominant period can be seen from a raw white noise signal (Figure 2A) and more clearly from the almost flat landscape on the power spectrum (Figure 2B). Applying to raw white noise signal a moving average filter of size 5-times the signal’s sampling interval (*M* = 5Δ*t*) leads to a smoothed noise signal (Figure 2C) that now has multiple dominant periods, as seen by the emergence of high power islands in Figure 2D. Comparing the original spectrum (Figure 2B) with the white noise smoothed spectrum (Figure 2D), it becomes evident that smoothing introduces a strong increase in Wavelet power for longer periods. In other words, smoothing perturbs the original signal by creating multiple high-power islands of long periods, also referred as spurious oscillations. To better capture the statistics behind these smoothing-induced spurious oscillations, it is best to look at the time averaged Wavelet spectrum. Figure 2E shows the mean expected power after smoothing white noise with a moving average filter. A zone of attenuated small periods become visible, the sloppy stopband for periods around ≲ 7Δ*t*. These are the fast, high-frequency components which get removed from the signal. However, for larger periods ≳ 10 the Wavelet power gets amplified up to 5-fold. It is this gain, given by 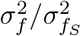, which leads to spurious results in the analysis.

As stated before, variance normalized white noise has a mean power of 1 for all frequencies or periods 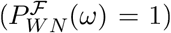. This allows to use a straightforward numerical method to estimate a filter response 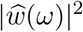, i.e. applying the smoothing operation to simulated white noise and time averaging the Wavelet spectra. This Monte Carlo approach works for every (also non-convolutional) smoothing method. Results for the Savitzky-Golay filter applied to white noise signals can be found in Supplementary Figure S2. Convolutional filters will in general produce more gain and hence more spurious oscillations with increasing window sizes in the time domain.

If smoothing, even with a rather small window (*M* = 5Δ*t*), already potentially introduces false positive oscillations, what does that mean for practical time-frequency analysis? For Wavelet analysis the answer is plain and clear: smoothing is simply not needed at all. A close inspection of the unaltered white noise Wavelet spectrum shown in Figure 2B, shows the same structures for higher periods as in the spectrum of the smoothed signal (Figure 2D). The big difference is, that even though these random apparent oscillations get picked up by the Wavelets, their low power directly indicates their low significance. As Wavelet analysis (see previous section) is based on convolutions, it already has power preserving *smoothing built in*. As illustration, we show in Figure 2F a raw noisy signal with lengthening period (noisy chirp) and the corresponding power spectrum (Figure 2F lower panel). Without any smoothing the main periodic signal can be clearly identified in the power spectrum. Thus, Wavelet analysis does not require smoothing for the detection of oscillations in very noisy signals. For all other spectrum analysis methods which rely on explicit smoothing, characteristics of the background noise and the signal to noise ratio are crucial to avoid detecting spurious oscillations. These are usually both quantities not readily available *a priori* or in practice.

### Detrending

Complementary to smoothing, an operation which removes the slow, low frequency components of a signal is generally called detrending. Strong trends can dominate a signal by effectively carrying most of the variance and power. There are at least two broad classes of detrending techniques: parametric fitting and convolution based. Both aim to estimate the trend as a function over time to be subtracted from the original signal. A parametric fit always is the best choice, if the deterministic processes leading to the trend are known and well understood. An example is the so called photobleaching encountered in time-lapse fluorescent imaging experiments, here an exponential trend can often be well fitted to the data based on first principle deliberations (Song et al. [1995]). However, there are often other slow processes, like cell viability or cells drifting in and out of focus, which usually can’t be readily described parametrically. For all these cases convolutional detrending with a window function *w*(*t*) is a good option and can be written as:

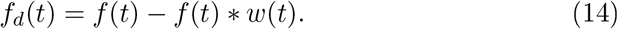

The trend here is nothing less than the smoothed original signal, i.e. *f*(*t*)∗*w*(*t*). However with the signal itself falling into the stop-band of the low-pass filter, with the aim to not capture and subtract any signal components.

Using basic algebra in the frequency domain we obtain an expression relating the window *w*(*t*) to the power spectrum of the original signal *f*(*t*):

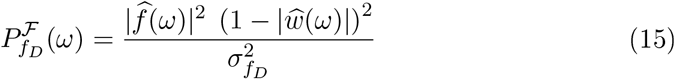

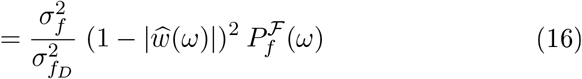

As in the case of smoothing, the so called *high-pass response* of the window function is given by 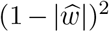 and scaled with the ratio of variances 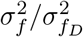. In strong contrast to smoothing, there is no overall gain in power in the range of the periods, passing through the filter (called *passband*). There is however, in case of moving average and other time-domain filters (see also Figure S2), no simple passband region. Instead, there are *rippling artifacts* in the frequency domain, meaning some periods getting amplified to up to 150% and others attenuated by up to 25%. To showcase why this can be problematic, we constructed a synthetic chirp signal sweeping through a range of periods *T*_1_ − *T*_2_, however, this time modified by a linear and an oscillatory trend (Figure 3A). The oscillatory component of the trend was chosen for clarity with a specific time scale given by its period *T_trend_*, which is three times the longest period found in the chirp signal. Strongly depending on the specific window size chosen for the moving average filter, there are various effects on both, the time and frequency domain (shaded area in Figure 3B) such as the introduction of amplitude envelopes and/or incomplete trend removal (Figure 3C). A larger window size is better to reduce the effect of the ripples inside the passband. However, the filter decay (*roll-off*) towards larger periods becomes very slow. That in turn means that trends can not be fully eliminated. Smaller window sizes perform better in detrending, but their passband can be dominated by ripples (see also supplementary Figure S3). In practice, sticking to filters originally designed for the time-domain without having oscillatory signals in mind, can easily lead to biased results of a time-frequency analysis. However, given the moderate gains of the detrending filter response, there is a much smaller chance to mistakenly detect spurious oscillations compared to the case of smoothing.

**Figure 3:**
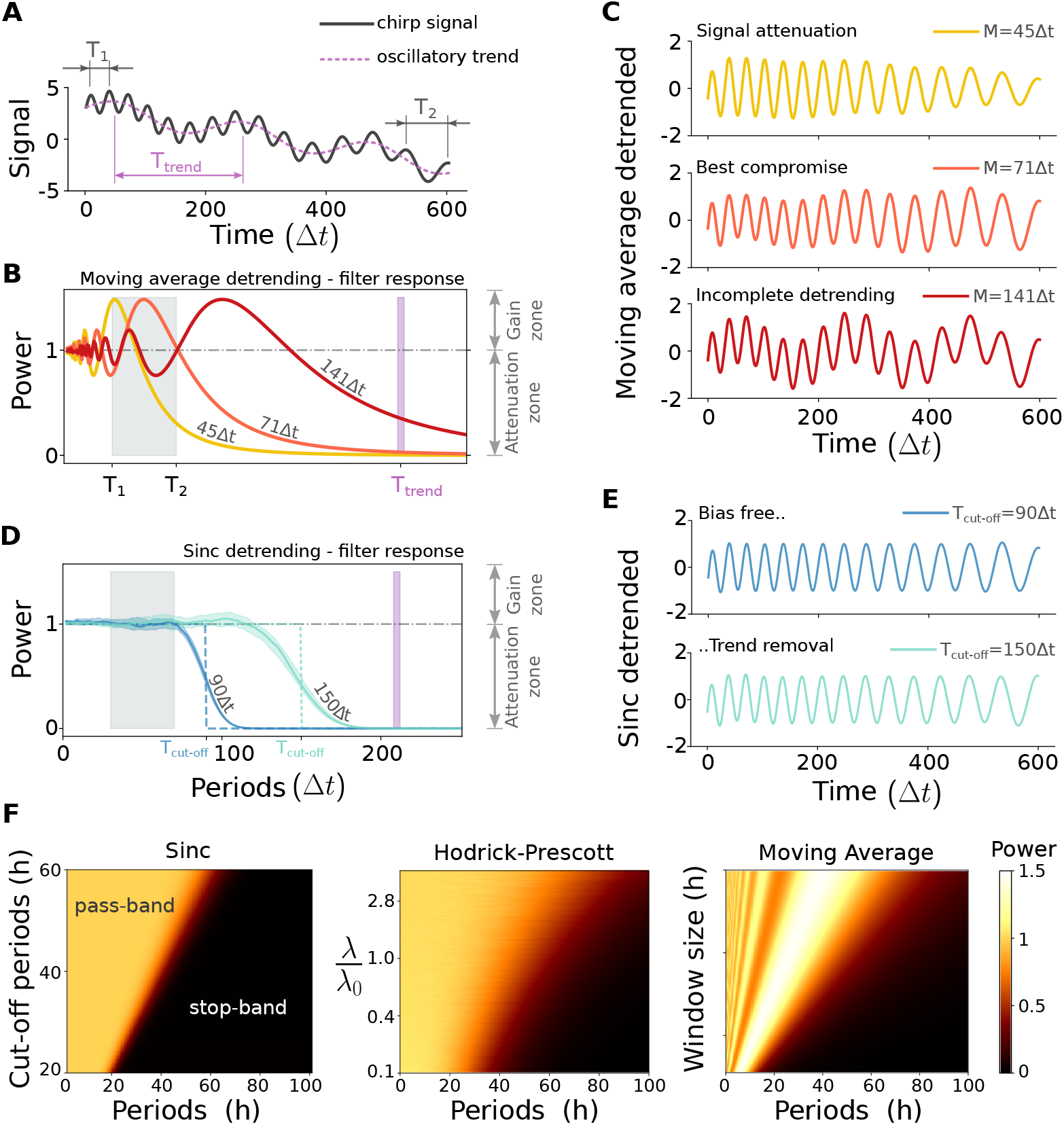
Detrending: moving average vs. optimal sinc. (A) Synthetic signal sweeping through the periods *T*_1_ = 30Δ*t* to *T*_2_ = 70Δ*t* within 600 sampling intervals (Δ*t*). This signal has an oscillatory and a linear trend: sin(2*π/T_trend_ t*) − 0.0066*t* + 1.5, with *T_trend_* = 210Δ*t*. (B) Detrending (or high-pass) response of the moving average filter with three different window sizes. The period range of the signal and the timescale of the trend are marked by shaded grey and violet regions respectively. (C) Time-domain effects of the moving average filters shown in B. A decreasing amplitude trend for *M* = 45Δ*t* (*top*), a best compromise case *M* = 71Δ*t*, and an incomplete trend removal for *M* = 141Δ*t* (*bottom*) can be observed. (D) Detrending response of the optimal sinc filter for two different cut-off periods as indicated. (E) Time-domain result for the sinc filters shown in C, indicating no amplitude effects while the trend is removed. (F) Parameter-dependent performance test of three common detrending filters, namely the sinc (*left*), Hodrick-Prescott (*middle*) and moving average (*right*), respectively. The sinc filter provides the clearest separation between pass- and stop-band. The reference value for the Hodrick-Prescott filter 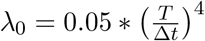 for *T* = 24Δ*t* is taken from Ravn and Uhlig [2002].

### The Sinc filter

The Sinc filter, also known as the optimal filter in the frequency domain, is a function with a constant value of one in the passband. In other words, frequencies which pass through are neither amplified nor attenuated. Accordingly, this filter also should be constantly zero in the stop-band, the frequencies (or periods) which should be filtered out. This optimal low-pass response can be formulated in the frequency domain simply as:

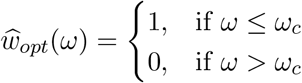

Here *ω_c_* is the *cut off frequency*, an infinitely sharp transition band dividing the frequency range into pass- and stop-band. It is effectively a box in the frequency domain (dashed lines in Figure 3D). Note that the optimal high-pass or detrending response simply and exactly swaps the pass- and stop-band. In the time domain via the inverse Fourier transform, this can be written as:

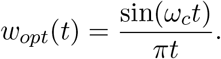

This function is known as the *sinc* function and hence the name *sinc filter*. An alternative name used in electrical engineering is brick-wall filter. In practice, this optimal filter has a nonzero roll-off as shown for two different cut-off periods 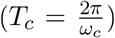 in Figure 3D. The sinc function mathematically requires the signal to be of infinite length. Therefore, every practical implementation implements *windowed sinc filters* (Smith et al. [1997]), see also supplementary information S3) about possible implementations. Strikingly still, there are no ripples or other artifacts in the frequency-response of the windowed sinc filter. And hence also the ‘real world’ version allows for a bias free time-frequency analysis. As shown in Figure 3E, the original signal can be exactly recovered via detrending. To showcase the performance of the sinc filter, we numerically compared its performance against two other common methods, the Hodrick-Prescott and moving average filter (Figure 3F). The stop- and passband separation of the sinc filter clearly is the best, although the Hodrick-Prescott filter with a parameterization as given by Ravn and Uhlig [2002] also gives acceptable results (see also supplementary Figure S5). The moving average is generally inadvisable, due to its amplification right at the start of the passband.

In addition to its advantages in practical signal analysis, the Sinc filter also allows to analytically calculate the gains from filtering pure noise (see also supplementary information A.3). The gain, and therefore the probability to detect spurious oscillations, introduced from smoothing is typically much larger compared to detrending. However, if a lot of energy of the noise is concentrated in the slow low frequency bands, also detrending with small cut-off periods alone can yield substantial gains (see Figure S6 and Figure S7). Importantly, when using the Sinc filter, the background spectrum of the noise will always be uniformly scaled by a constant factor in the pass-band. There is no mixing of attenuation and amplification as for time-domain filters like moving average (Figure 3B and C). If the spectrum of the noise can be estimated, or an empirical background spectrum is available, the theory presented in A.3 allows to directly calculate the correct confidence intervals.

## 4 Readout - Along the Ridge

Wavelet power spectra are by itself already a valuable output of a signal analysis and give a direct visual representation of the relative weight of periods (frequencies) present in the signal of interest in a time dependent manner. However, especially in the context of biological rhythms, there often is the well justified assumption that there is only one main oscillatory component present ^1^. The extraction of the instantaneous period, amplitude and phase of that component is of prime interest for the practitioner. In this section we show how to obtain these important features using wavelet transforms as implemented in `pyBOAT`.

### Tracking the main oscillatory components

From the perspective of Wavelet power spectra, such main oscillatory components are characterized by concentrated and time-connected regions of high power. *Wavelet ridges* are a means to trace these regions on the time-period plane. For the vast majority of practical applications a simple maximum ridge extraction is sufficient. This maximum ridge can be defined as:

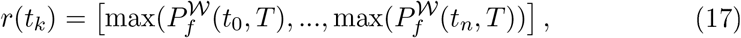

with *n* being the number of sample points in the signal. Thus, the ridge *r*(*t_k_*) maps every time point *t_k_* to a row of the power spectrum, and therewith to a specific instantaneous period *T_k_* (Figure 4C and D). Evaluating the power spectrum along a ridge, gives a time series of powers: 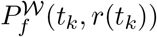. Setting a power threshold value is recommended to avoid evaluating the ridge in regions of the spectrum where the noise dominates (In Figure 4C threshold is set to 5). Alternatively to simple maximum ridge detection, more elaborated strategies for ridge extraction have been proposed (Carmona et al. [1995]).

**Figure 4:**
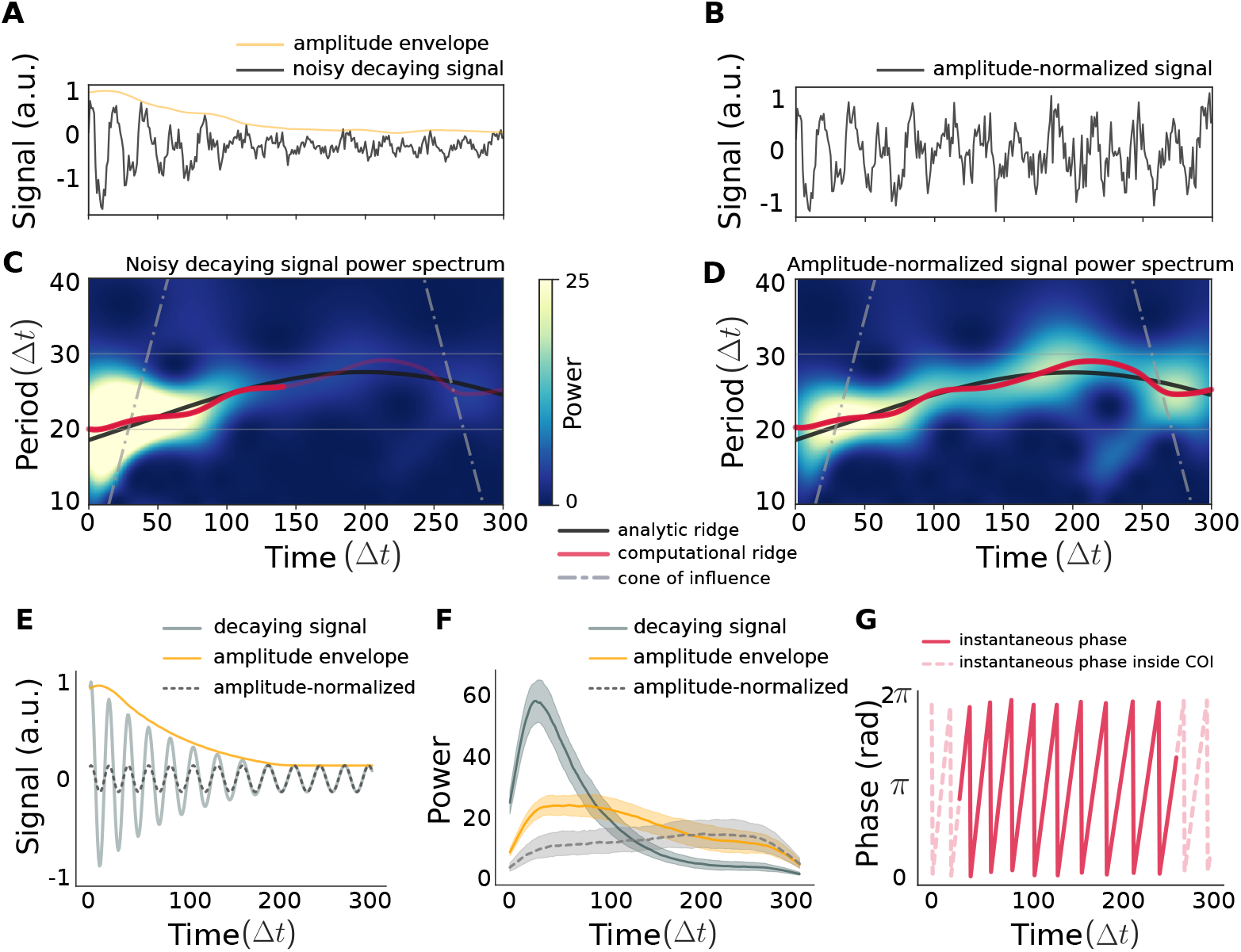
Ridge detection and amplitude normalization. (A) A noisy synthetic signal with a decaying amplitude envelope (orange line). (B) Signal from panel A after amplitude normalization. (C) Wavelet power spectrum of the signal in panel A. The decaying amplitudes lead to very low powers towards the end of the signal. The extracted maximum power ridge (red line) reaches the lower threshold of 5 in the middle part of the signal (indicated by the light red line transition). As a reference, the (expected) analytic ridge is depicted by a black line. The cone of influence is indicated by gray dashed lines. (D) Wavelet power spectrum of the normalized signal in panel B. The oscillations are well captured, indicated by the high power during at all times. Hence, the thresholded maximum ridge traces the whole signal. (E) The deterministic parts of the synthetic signal shown in A. (F) Median and quartiles of the power along non-thresholded ridges of the signals as indicated (*N_trial_* = 50). Signals with amplitude decay have only very low powers towards the end. The amplitude normalized signals do not have a constant signal-to-noise ratio even after normalization, and therefore still have higher power on average in the beginning as compared to the signal without envelope. (G) Instantaneous phase extracted from the amplitude normalized signal. The COI extent is marked by dashed lines.

### Amplitude normalization

A problem often encountered when dealing with biological data is a general time-dependent amplitude envelope (Figure 4A). Under our wavelet approach, the power spectrum is normalized with the overall variance of the signal. Consequently, regions with low signal amplitudes but robust oscillations are nevertheless represented as very low power blurring them with the spectral floor (Figure 4C). This leads to the impractical situation, where even a noise free signal with an amplitude decay will show very low power at the end (Figure 4E,F and S8), defeating its statistical purpose. A practical solution in this case is to estimate an amplitude envelope and subsequently normalize the signal with this envelope (Figure 4A and B). We specifically show here non-periodic envelopes, estimated by a sliding window (see also methods). After normalization, lower amplitudes are no longer penalized and an effective power-thresholding of the ridge is possible (Figure 4D and F).

A limitation of convolutional methods, including Wavelet-based approaches, are *edge effects*. At the edges of the signal, the Wavelets only partially overlap with the signal leading to a so-called *cone of influence* (COI) (Figure 4C and D). Even though the periods are still close to the actual values, phases and especially the power should not be trusted inside the COI (see discussion and supplementary Figure S9).

Once the trace of consecutive Wavelet power maxima has been determined and thresholded, evaluating the transform along it yields the instantaneous envelope amplitude, normalized amplitude and phases (see Figure 4E,F and G). To obtain the amplitudes, we derive a simple scaling rule for the power time series to be converted into amplitudes (see derivations in A.4).

### Applications

After introducing the different time series analysis steps using synthetic data for clarity, in this paragraph, we discuss examples of `pyBOAT` applications to real data. To showcase the versatility of our approach, we chose datasets obtained from different scientific fields.

In Figure 5A we display COVID-19 infections in Italy as reported by the Disease Prevention and Control (ECDC). A Sinc-filter trend identification with a cut-off period of 14 days reveals a steep increase in newly reported infections at the beginning of March and a steady decline after the beginning of April. Subtracting this non-linear trend clearly exposes oscillations with a stable period of one week (Figure 5B, and see Supplementary Figure S10A for power spectrum analysis). Similar findings were recently reported in Ricon-Becker et al. [2020].

**Figure 5:**
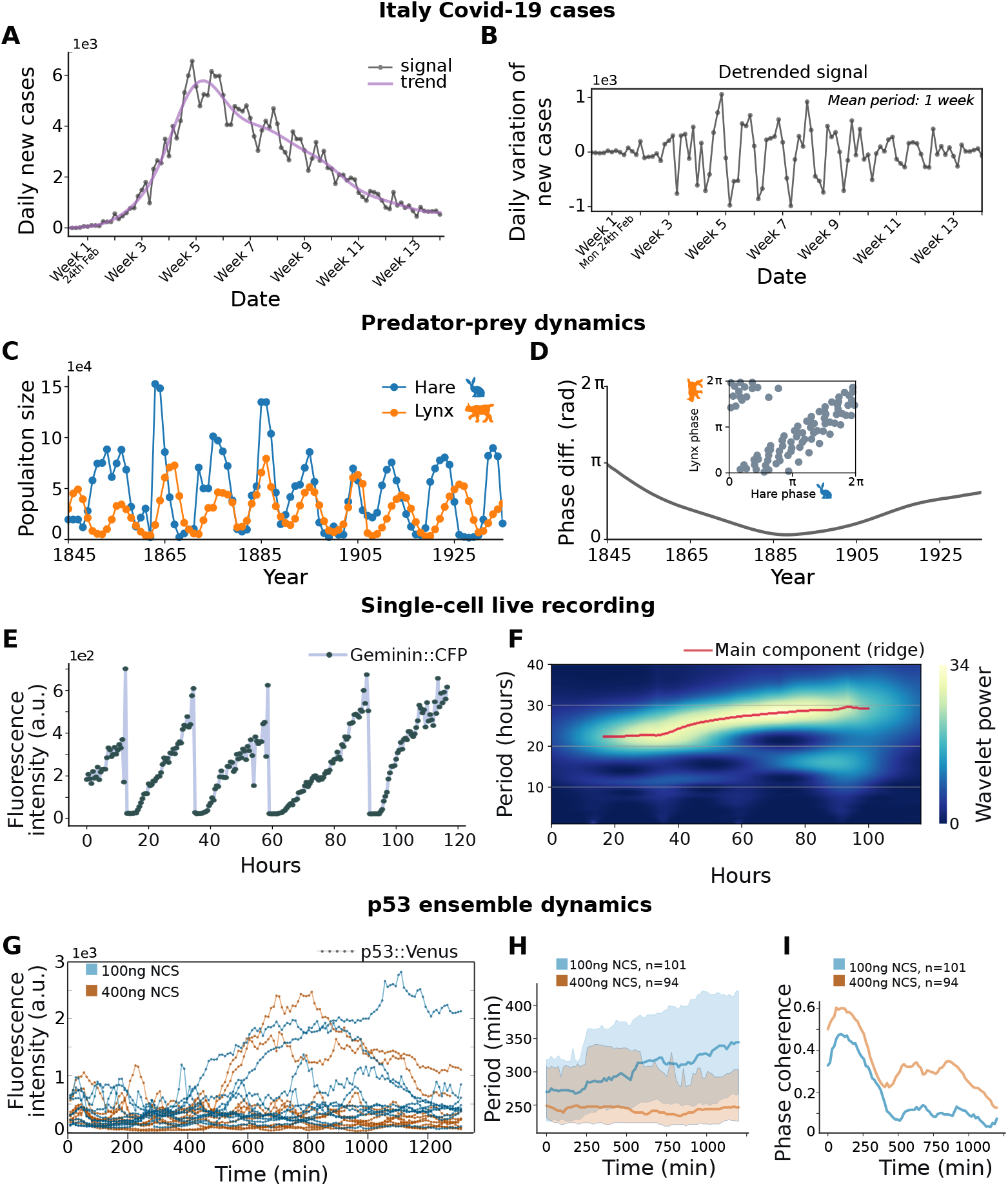
Time-dependent analysis of oscillations using pyBOAT. A) Daily confirmed COVID-19 infections in Italy (gray line; dots represent single data points) from 19th Feb to May 25th 2020. Non-linear baseline trend as determined by the Sinc-filter, using a cut-off period of 14 days (purple line). B) Daily oscillatory variation of Covid-19 new cases. Plot in B) was obtained by removing the trend from the data on panel A. C) Population number of snowshoe hares (blue) and lynx (orange). D) Difference between the instantaneous (i.e. time-dependent) phases of the hare and lynx oscillations as extracted from the wavelet-spectra ridges. In the inset, instantaneous phases of the hare and the lynx are plotted against each other. E) Single-cell trajectory of a U2OS cell carrying a Geminin-CFP fluorescent reporter. F) Wavelet analysis reveals a strong main periodic component (ridge in red). G) Single-cell trajectories of MCF7 cells carrying a p53-venus fluorescent reporter under low 100ng NCS and high 400ng NCS DNA damage. 10 randomly selected traces by condition. H) Median and quartiles of the detected periods over time. I) Phase-coherence over time as a measure for synchronicity for each condition.

The signals shown in Figure 5C show cycles in Hare-Lynx population sizes as inferred from the yearly number of pelts, trapped by the Hudson Bay Company. Data has been taken from Odum and Barrett [1971]. The corresponding power spectra are shown in the supplement (Figure S10B), and reveal a fairly stable 10 year periodicity. After extracting the instantaneous phases with `pyBOAT`, we calculated the time-dependent phase differences as shown in Figure 5B. Interestingly, the phase difference slowly varies between being almost perfectly out of phase and being in phase for a few years around 1885.

The next example signal shows a single-cell trajectory of a U2OS cell carrying a Geminin-CFP fluorescent reporter (Granada et al. [2020]). Geminin is a cell cycle reporter, accumulating in the G2 phase and then steeply declining during mitosis. Applying `pyBOAT` on this non-sinusoidal oscillations reveals the cell-cycle length over time (Figure 5F), showing a slowing down of the cell cycle progression for this cell. Ensemble dynamics for a control and a cisplatin treated population are shown in the Supplementary Figure S10C.

The final example data set is taken from Mönke et al. [2017], here populations of MCF7 cells where treated with different dosages of the DNA damaging agent NCS. This in turn elicits a dose-dependent and heterogeneous p53 response, tracked in the individual cells for each condition (Figure 5G). `pyBOAT` also features readouts of the ensemble dynamics: Figure 5H shows the time-dependent period distribution in each population, Figure 5I the phase coherence over time. The latter is calculated as 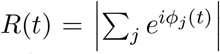. It ranges from zero to one and is a classical measure of synchronicity in an ensemble of oscillators Kuramoto [1984]. The strongly stimulated cells (400ng NCS) show stable oscillations with a period of around 240min, and retain more phase coherent after an initial drop in synchronicity. The medium stimulated cells (100ng NCS) start to slow down on average already after the first pulse, both the spread of the period distribution and the low phase coherence indicate a much more heterogeneous response. Two individual cells and their wavelet analysis are shown in Supplementary Figure S10D.

### Graphical interface

The extraction of period, amplitude and phase is the final step of our proposed analysis workflow, which is outlined in the screen-captures of Figure 6. The user interface is separated into several sections. First, the ‘DataViewer’ allows to visualize the individual raw signals, the trend determined by the Sinc filter, the detrended time series and the amplitude envelope. Once satisfactory parameters have been found, the actual Wavelet transform together with the ridge are shown in the ‘Wavelet Spectrum’ window. After ridge extraction, it is possible to plot the instantaneous observables in a ‘Readout’ window. Each plot produced from the interface can be panned, zoomed in and saved separately if needed.

**Figure 6:**
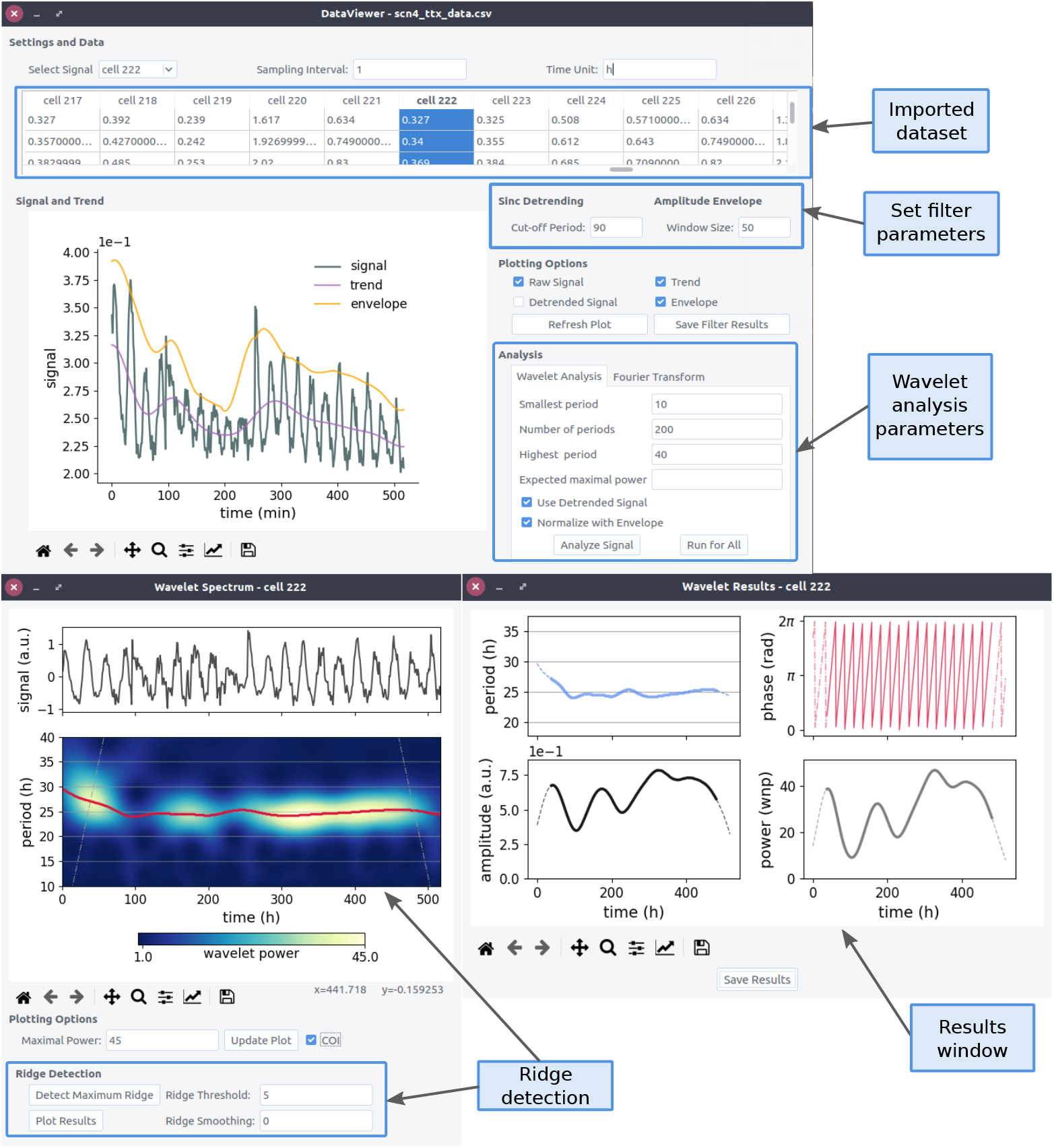
Screen captures of the elements of pyBOAT in action. The signal gets detrended with a cut-off period of 90h, and an amplitude envelope is estimated via a window of size 50h. See labels and main text for further explanations. The example trajectory displays a circadian rhythm of 24h and is taken from the data set published in Abel et al. [2016].

Once the fidelity of the analysis has been checked for individual signals, it is also possible to run the entire analysis as a ‘batch process’ for all signals imported. One aggregated result which we found quite useful is to determine the ‘rhythmicity’ of a population by creating a histogram of time-averaged powers of the individual ridges. A classification of signals into ‘oscillatory’ and ‘non-oscillatory’ based on this distribution, e.g. by using classical thresholds (Otsu [1979]) is a potential application. Examples of the provided ensemble readouts are shown in Figure 5H and I and Supplementary Figure S10C.

Finally, `pyBOAT` also features a synthetic signal generator, allowing to quickly explore its capabilities also without having a suitable dataset at hand. A synthetic signal can be composed of up to two different chirps, and AR1-noise and an exponential envelope can be added to simulate possible challenges often present in real data (see also Material and Methods and Supplementary Figure S11).

Installation and user guidelines of `pyBOAT` can be found in the github repository.

## 5 Discussion

Recordings of biological oscillatory signals can be conceptualized as an aggregate of multiple components, those coming from the underlying system of interest and additional confounding factors such as noise, modulations and trends that can disguise the underlying oscillations. In cases of variable period with noisy amplitude modulation and non-stationary trends the detection and analysis of oscillatory processes is a non-trivial endeavour. Here we introduced `pyBOAT`, a novel software package that uses a statistically rigorous method to handle non-stationary rhythmic data. `pyBOAT` integrates pre- and post-processing steps without making *a priori* assumptions about the sources of noise and periodicity of the underlying oscillations. We showed how the signal processing steps of smoothing, detrending, amplitude envelope removal, signal detection and spectral analysis can be resolved by our hands-off standalone software (Figure 1 and 5).

Artifacts introduced by the time series analysis methods itself are a common problem that inadvertently disturbs results of the time-frequency analysis of periodic components (Wilden et al. [1998]). Here we first analyzed the effects of data-smoothing on a rhythmic noisy signal and showed how common smoothing approaches disturb the original recordings by introducing non-linear attenuations and gains to the signal (Figures 2, S6 and S7). These gains easily lead to spurious oscillations that were not present in the original raw data. These artifacts have been characterized since long for the commonly used moving-average smoothing method, known as the Slutzky-Yule effect (Slutzky [1937]). Using an analytical framework, we describe the smoothing process as a filter operation in frequency domain. This allows us to quantify and directly compare the effects of diverse smoothing methods by means of response curves. Importantly, we show here how any choice of smoothing unavoidably transforms the original signal in a non-trivial manner. One potential reason for its prevalence is that practitioners often implement a smoothing algorithm without quantitatively comparing the spectral components before versus after smoothing. `pyBOAT` avoids this problem by implementing a wavelet-based approach that *per se* evades the need to smooth the signal.

Another source of artifacts are detrending operations. Thus, we next studied the spectral effects that signal detrending has on rhythmic components. Our analytical and numerical approaches allowed us to compare the spectral effects of different detrending methods in terms of their response curves (see Figure 3). Our results show that detrending also introduces non-trivial boosts and attenuations to the oscillatory components of the signal, strongly depending on the background noise (Figures S6 and S7). In general there is no universal approach and optimally a detrending model is based on information about the sources generating the trend. In cases without prior information to formulate a parametric detrending in the time domain, we suggest that the safest method is the convolution based *sinc filter*, as it is an “ideal” (step-function) filter in the frequency domain (Figures 3C and S3). Furthermore we compared the performance of the sinc filter with two other commonly applied methods to remove non-linear trends in data (Figure 3F), i.e. the moving average (Díez-Noguera [2013]) and Hodrick-Prescott (Myung et al. [2012], Schmal et al. [2018], St. John and Doyle [2015]) filter.

In addition to smoothing and detrending, amplitude-normalization by means of the amplitude envelope removal is another commonly used data processing step that `pyBOAT` is able to perform. Here we further show how that for decaying signals amplitude normalization grants that the main oscillatory component of interest can be properly identified in the power spectrum (Figure 4A to D). This main component is identified by a ridge-tracking approach that can be then used to extract instantaneous signal parameters such as amplitudes, power and phase (Figure 4E to G).

Rhythmic time series can be categorized into those showing stationary oscillatory properties and the non-stationary ones where periods, amplitudes and phases change over time. Many currently available tools for the analysis of biological rhythms rely on methods aimed at stationary oscillatory data, using either a standalone software environment such as `BRASS` (Edwards et al. [2010], Locke et al. [2005]), `ChronoStar` (Klemz, et al. [2017]) and `CIRCADA` (Cenek et al. [2020]) or an online interfaces such as `BioDare` (Moore et al. [2014], Zielinski et al. [2014]).

Continuous wavelet analysis allows to reveal non-stationary period, amplitude and phase dynamics and to identify multiple frequency components across different scales within a single oscillatory signal (Leise [2013], Leise et al. [2012], Rojas et al. [2019]) and is thus complementary to approaches that are designed to analyze stationary data. In contrast to the `R`-based `waveclock` package (Price et al. [2008]), `pyBOAT` can be operated as a standalone software tool that requires no prior programming knowledge as it can be fully operated using its graphical user interface (GUI). An integrated batch processing option allows the analysis of large data sets within a few “clicks”. For the programming interested user, `pyBOAT` can also be easily scripted without using the GUI, making it simple to integrate it into individual analysis pipelines. `pyBOAT` also distinguishes itself from other wavelet-based packages (e.g. Harang et al. [2012]) by adding a robust *sinc filter*-based detrending and a statistically rigorous framework, providing the interpretation of results by statistical confidence considerations.

`pyBOAT` is not specifically designed to analyze oscillations in high-throughput “omics” data. For such sake, specialized algorithms such as ARSER (Yang and Su [2010]), JTK-cycle (Hughes et al. [2010]), MetaCycle (Wu et al. [2016]) or RAIN (Thaben and Westermark [2014]) are more appropriate. Its analysis reveals basic oscillatory properties such as the time-dependent (instantaneous) rhythmicity, period, amplitude and phase but is not aimed at more specific statistical tests such as, e.g., tests for differential rhythmicity as implemented in DODR (Thaben and Westermark, 2016). The continous-wavelet analysis underlying `pyBOAT` requires equidistant time series sampling with no gaps. Methods such as Lomb-Scargle periodograms or harmonic regressions are more robust or even specifically designed with respect to unevenly-sampled data (Lomb [1976], Ruf [1999]). Being beyond the scope of this manuscript, it will be interesting in future work to integrate the ability to analyze unevenly sampled data into the pyBOAT software, either by the imputation of missing values (e.g. by linear interpolation) or the usage of wavelet functions specifically designed for this purpose (Thiebaut and Roques [2005]).

`pyBOAT` is fast, easy too use and statistically robust analysis routine designed to complement existing methods and advance the efficient time series analysis of biological rhythms research. In order to make it publicly available, `pyBOAT` is a free and open-source, multi-platform software based on the popular Python (Van Rossum and Drake [2009]) programming language. It can be downloaded using the following link: https://github.com/tensionhead/pyBOAT, and is available on the Anaconda distribution (via the *conda-forge* channel).

## 6 Methods

### Software

`pyBOAT` is written in the Python programming language (Van Rossum and Drake [2009]). It makes extensive use of Python’s core scientific libraries *numpy* and *scipy* (Virtanen et al. [2020]) for the numerics. Additionally we use *matplotlib* (Hunter [2007] for visualization, and *pandas* (McKinney [2010]) for data management. `pyBOAT` is released under the open source GPL-3.0 license, and its code is freely available from https://github.com/tensionhead/pyBOAT. The readme on this repository contains further information and installation instructions. `pyBOAT` is also hosted on the popular Anaconda distribution, as part of the *conda-forge* community https://conda-forge.org/.

### Amplitude envelope

To estimate the amplitude envelope in the time domain, we employ a moving window of size *L* and determine the minimum and maximum of the signal inside the window for each time point *t*. The amplitude at that time point is then given by 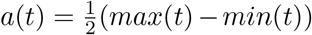. This works very well for envelopes with no periodic components, like an exponential decay. However, this simple method is not suited for oscillatory amplitude modulations. It is also recommended to sinc-detrend the signal before estimating the amplitude envelope. Note that *L* should always be larger then the maximal expected period in the signal, as otherwise the signal itself gets distorted.

### Synthetic signals

A noisy chirp signal can be written as: *f*(*t_i_*) = *a cos*(*ϕ*(*t_i_*)) + *d x*(*t_i_*), where *ϕ*(*t_i_*) is the instantaneous phase and the *x*(*t_i_*) are samples from a stationary stochastic process (the background noise). The increments of the *t_i_* are the sampling interval: *t_i_*_+1_ − *t_i_* = Δ*t*, with *i* = 0, 1*,…, N* samples. Starting from a linear sweep through angular frequencies: *ω*(0) = *ω*_1_ and *ω*(*t_N_*) = *ω*_2_, we have 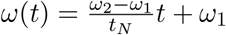. The instantaneous phase is then given by

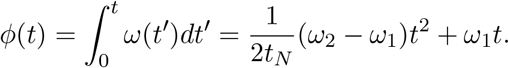

Sampling *N* times from a gaussian distribution with standard deviation equal to one corresponds to gaussian white noise *ξ*(*t_i_*). With *x*(*t_i_*) = *ξ*(*t_i_*) the signal to noise ration (SNR) then is *a*^2^*/d*^2^.

A realization of an AR1, process can be simulated by a simple generative procedure: the inital *x*(*t*_0_) is a sample from the standard normal distribution. Then the next sample is given by: *x*(*t_i_*) = *αx*(*t_i−_*_1_) + *ξ*(*t_i_*), with *α* < 1.

Simulating pink noise is less straightforward, and we use the Python package *colorednoise* from https://pypi.org/project/colorednoise for the simulations. Its implementation is based on Timmer and Koenig [1995].

## Acknowledgments

We gratefully thank Bharath Ananthasubramaniam, Hanspeter Herzel, Pedro Pablo Rojas and Shaon Chakrabarti for fruitful discussions and comments on the manuscript. We further thank Jelle Scholtalbers and the GBCS unit at the EMBL in Heidelberg for technical support. We thank members of the Aulehla and Leptin labs for comments, support and helpful advice.

## Footnotes

1 This is also the underlying assumption of the Hilbert transform, the main component being the so called ‘analytical signal’ (Hussain and Boashash [2002]).

