## Supplementary Material for "Optimal time frequency analysis for biological data - pyBOAT"

### A. Supplementary Information

#### A.1. General Wavelet Properties

Here we want to briefly discuss a few mathematical properties of Wavelets in general, formal introductions may for example be found in Daubechies [1992] and Mallat [1999]. An important property of Wavelets is the vanishing average:

$$\int_{-\infty}^{\infty} \Psi_{s,\tau}(t)\, dt = 0 \qquad (1)$$

which is often refered to as "admissibility criterion". Grossmann and Morlet have shown, that for a faithful decomposition and re-synthesis this condition has to be fullfilled (Grossmann and Morlet [1985]). In Fourier space this is equivalent to $\widehat{\Psi}_{s,\tau}(0) = 0$, meaning a vanishing zero-frequency component.

To ensure equal decomposition on all scales, a normalization to unit energy is required:

$$\int_{-\infty}^{\infty} |\Psi_{s,\tau}(t)|^2\, dt = 1 \qquad (2)$$

These two conditions alone imply that a Wavelet function $\Psi_{s,\tau}(t)$ is oscillatory and decaying in time.

##### A.1.1. Center frequency

Wavelets are localized in frequency space but in contrast to the basis functions of the Fourier transform, they do not have a single frequency. However, it is possible to still associate one single frequency to an individual Wavelet with scale $s$ by defining the *center frequency*:

$$\omega_{center}(s) = \int_{-\infty}^{\infty} \omega |\widehat{\Psi}_{s,\tau}(\omega)|^2\, d\omega \qquad (3)$$

From equation (2) and Plancherel's theorem it follows that the Fourier power spectrum of a Wavelet is also normalized: $\int |\widehat{\Psi}_{s,\tau}(\omega)|^2 d\omega = 1$, and therefore $\omega_{center}$ simply is the mean of its Fourier power spectral density.

### A.2. Morlet Wavelet Properties

Taking the usual definition (Torrence and Compo [1998]) of the Morlet mother wavelet:

$$\Psi(t) = \pi^{-1/4}\, e^{-t^2/2}\, e^{i\omega_0 t}, \tag{4}$$

its Fourier transform reads as

$$\widehat{\Psi}(\omega) = \sqrt{2\pi}\, e^{-\frac{1}{2}(\omega-\omega_0)^2} \tag{5}$$

which is a shifted Gaussian distribution with mean $\omega_0$ as the center frequency and unit standard deviation. From

$$\widehat{\Psi}(0) = \sqrt{2\pi} e^{-\frac{1}{2}\omega_0^2}, \tag{6}$$

we see that the admissibility condition (1) is not fullfilled. In the literature this form of the Morlet wavelet is still the most prominent due to it's simplicity. To at least approximately comply with the admissibility criterion, an $\omega_0 \geq 2\pi$ is often chosen such that $\widehat{\Psi}(0)$ is close to zero as

$$\widehat{\Psi}(0, \omega_0 = 2\pi) = \sqrt{2\pi} e^{-\frac{1}{2}4\pi^2} \approx 3.78 \cdot 10^{-9}. \tag{7}$$

From this we find that the center frequency of the Morlet wavelet with scale $s$ reads as $\omega_c(s) = \frac{\omega_0}{s}$, using equations (3) and (5).

However, Torrence and Compo [1998] give the Morlet center frequency as

$$\omega_c = \frac{s}{2}(\omega_0 + \sqrt{2 + \omega_0^2}) \tag{8}$$

which is inconsistent with the definition of equation 4. Equation (A.1.1) stems from the exactly admissible form of the Morlet wavelet Ashmead [2010]

$$\Psi_{adm}(t) \sim e^{\frac{-t^2}{2}} \left( e^{i\omega_0 t} - e^{-\frac{1}{2}\omega_0^2} \right), \tag{9}$$

and its Fourier transform is given by

$$\widehat{\Psi}_{adm}(\omega) \sim e^{-\frac{1}{2}(\omega-\omega_0)^2} - e^{-\frac{1}{2}(\omega_0^2+\omega^2)}, \tag{10}$$

which readily fulfills the admissibility criterion $\widehat{\Psi}_{adm}(0) = 0$. For $\omega_0 = 2\pi$ the deviation between the two center frequency definitions is around 1.25%, with the latter one being more exact.

### A.3. Calculating gains

As shown in section 3 of the main text, gains in the Fourier (or time averaged Wavelet) spectra stem from the ratio of variances before and after convolutional filtering with a window function $w(t)$. We define the gain from smoothing as

$$\gamma_S = \frac{\sigma_f^2}{\sigma_{f_S}^2}, \tag{11}$$

where $\sigma_f^2$ is the variance of the unfiltered signal, and $\sigma_{f_S}^2$ is the variance of the filtered signal. Similarly we define the gain from detrending as

$$\gamma_D = \frac{\sigma_f^2}{\sigma_{f_D}^2}, \tag{12}$$

where $\sigma_{f_D}^2$ is the variance after detrending. Note that $\int_{-\infty}^{+\infty} w(t)dt = 1$, $\gamma_S \geq 1$ and $\gamma_D \geq 1$ holds in general, since the window functions are normalized. This means that the gains can be attributed to a *loss of variance* in the filtered signals.

Our general approach to calculate this loss is to identify the variance of a signal, $\sigma_f^2 = Var(f)$, with the *energy* contained in it's Fourier power spectrum $P_f(\omega) = |\widehat{f}(\omega)|^2$. For $f(t)$ being a zero-mean stationary process, we can write Plancherel's theorem as:

$$Var(f) = \int_0^\infty P_f(\omega)d\omega, \tag{13}$$

such that we get

$$Var(w * f) = \int_0^\infty |\widehat{w}(\omega)|^2 P_f(\omega)d\omega, \tag{14}$$

for a convolutional filtering, where $\widehat{w}(\omega)$ is the Fourier transform of the window.

The above integral for the variance of a filtered signal holds for arbitrary window functions. Now we employ the sinc filter which, as defined in the main text, is a box extending from zero to the cut-off frequency $\omega_c$ in the frequency domain. Thus, application of the sinc filter solely changes the limits of integration:

$$Var(w_{sinc} * f) = \int_0^{\omega_c} P_f(\omega)d\omega. \tag{15}$$

This expression is the integrated power spectral density of the original unfiltered signal from zero to the cut-off frequency $\omega_c = 2\pi/T_c$. For detrending we similarly obtain:

$$Var(f - w_{sinc} * f) = \int_0^\infty P_f(\omega)d\omega - \int_0^{\omega_c} P_f(\omega)d\omega \tag{16}$$

$$= \int_{\omega_c}^\infty P_f(\omega)d\omega \tag{17}$$

Experimentally obtained signals are usually sampled at a given interval $\Delta t$, such that the upper limit for the frequencies is given by the Nyquist frequency $2\pi f_{Ny} = \omega_{Ny} = \pi/\Delta t$. The lower bound for the integration is given by $1/N\Delta t$ which approaches zero for very long signals. Using the definitions

$$I(\omega) = \int_0^\omega P_f(\omega')d\omega', \tag{18}$$

$$S = I(\omega_c)\, and$$

$$D = I_{Ny} - I(\omega_c)$$

where $I_{Ny} \equiv I(\omega_{Ny}) - I(0)$ is the *total variance* up to the Nyquist frequency (such that we have $I_{Ny} = \sigma_f^2$), $S = \sigma_{f_S}^2$ is the variance after sinc-smoothing, and $D = \sigma_{f_D}^2$ is the variance left after sinc-detrending, we can eventually obtain the gains

$$\gamma_S = \frac{I_{Ny}}{S} \; and \qquad (19)$$
$$\gamma_D = \frac{I_{Ny}}{D} = \frac{I_{Ny}}{I_{Ny} - S},$$

with $I_{Ny} = S + D$.

To calculate the gains for a specific null model (i.e., the background noise spectrum), we simply have to integrate the respective power spectral densities in the limits defined above. In the following we investigate three common noise models and calculate their respective spectral gains attained from sinc-filtering as a function of the cut-off period $T_c = 2\pi/\omega_c$. The results are summarized in Figure S6 and Figure S7. Note that Plancherel's theorem depends on the *frequency representation* of a given spectrum. The frequency is the physical quantity related to the energy contained in a spectrum.

To compare our findings to simulations, the respective noise process was filtered with a sinc-filter with a cut-off period of $T_c = 10\Delta t$. We then time-averaged the Wavelet power spectra of 50 realizations and plotted the numerical against the analytical results:

$$\mathcal{P}_{f_S}(\omega) = \gamma_S(T_c) \times \mathcal{P}_f(\omega) \qquad (20)$$
$$\mathcal{P}_{f_D}(\omega) = \gamma_D(T_c) \times \mathcal{P}_f(\omega).$$

As predicted, rescaling the unfiltered power spectra with the calculated gains gives the observed power spectra (see Figure S7). In principle this allows to adapt the confidence levels as outlined in the main text and in reference Torrence and Compo [1998] to accomodate for sinc-filtering. However, in practice this requires to first estimate the noise spectrum that is present in the data and subsequently solve for the integrated power spectrum.

#### A.3.1. White noise

White noise is the simplest null model with a flat spectrum that is equal to one for all frequencies. Specifically Gaussian white noise can be written as

$$f(t_i) = \xi(t_i), \text{ with } \xi \sim \mathcal{N}(0, \sigma^2)\,. \qquad (21)$$

It is the most random of all stochastic processes, indicated by it's vanishing autocorrelation $acorr(f_{WN}) = \delta(\tau)$, where $\delta(\tau)$ denotes Dirac's delta function.

With $\mathcal{P}_f(\omega) = 1$, the integral expressions defined in equations 18 and 19 reduce to simple algebra, and the gains read as

$$\gamma_S(T_c) = \frac{1}{2} T_c \tag{22}$$

$$\gamma_D(T_c) = \frac{T_c}{T_c - 2}\,. \tag{23}$$

The decomposition of the power spectrum into $S$ and $D$ for $T_c = 10\Delta t$ and the solution for the gains is shown in Figure S6. The numerical results for $T_c = 10\Delta t$ coincide perfectly with the analytical results, compare Figure S7.

#### A.3.2. Autoregressive model AR1

A stochastic process incorporating actual correlations in time is given by:

$$f(t_i) = \alpha f(t_{i-1}) + \xi(t_i), \tag{24}$$

with $\alpha < 1$ and $\xi(t_i)$ being white noise. The corresponding power spectrum is given as:

$$\mathcal{P}_f(\omega, \alpha) = \frac{1 - \alpha^2}{1 + \alpha^2 - 2\alpha \cos(\omega)} \tag{25}$$

To evaluate the gain, we need the integrated power spectral density:

$$I(\omega) = -\frac{1}{\pi} \arctan\left(\frac{\alpha + 1}{\alpha - 1} \tan(\omega/2)\right) \tag{26}$$

Using equations (18) and (19) readily provides the gains $\gamma_S(T_c, \alpha)$ and $\gamma_D(T_c, \alpha)$ as a function of the AR1-paramter $\alpha$ and the cut-off period $T_c$. Figure S6 (left and middle panel) depicts the AR1 power spectrum and its decomposition in $S$ and $D$ for $\alpha = 0.4$ and $T_c = 10\Delta t$.

Smaller values for $\alpha$ will tip the gains more towards white noise behavior, whereas a larger $\alpha$ will move the intersection point of the gain functions towards higher cut-off periods (Figure S6 right panel). Again, the numerical gains coincide precisely with the analytic results for $\gamma_S(T_c, \alpha)$ and $\gamma_D(T_c, \alpha)$ (Figure S7).

#### A.3.3. Pink Noise

The power spectrum of a $1/f^\beta$ noise process, with $\omega = 2\pi f$, reads as

$$\mathcal{P}_f(\omega) = \frac{1 - \beta}{\pi^{1-\beta}} \omega^{-\beta} \tag{27}$$

leading to

$$I(\omega) = \left(\frac{\omega}{\pi}\right)^{1-\beta}, \tag{28}$$

where $\beta < 1$ is enforced to ensure $I(0) = 0$. Again, by using equations (18) and (19) the gains read

$$\gamma_S(T_c, \beta) = \left(\frac{T_c}{2}\right)^{1-\beta} \tag{29}$$

$$\gamma_D(T_c, \beta) = \frac{1}{1 - (2/T_c)^{1-\beta}} \, . \tag{30}$$

We plot the decomposition of the power spectrum for $\beta = 0.7$ and $T_c = 10\Delta t$ in Figure S6 (left and middle panel). We additionally plotted two sets of gain functions for $\beta = 0.4$ and $\beta = 0.7$ (Figure S6 right panel). Smaller values of $\beta$ lead to more power in the higher frequency region of the spectrum, and hence larger (smaller) gains from smoothing (detrending).

#### A.3.4. Evaluation of the results

Even though the analytic calculations where done exclusively for the sinc-filter, the results easily generalise: It is the ratio of the variance left after smoothing ($S$) compared to what is left after detrending ($D$) which determines the bias introduced into the analysis. Here, not the characteristics of the filter itself, but the chosen (or measured) null model for the background noise and its energy distribution on the frequency axis are the determining factors. Additionallly, the ratio of the sampling interval $\Delta t$ to the cut-off period (in measurement units) is key for using our results. Detrending a signal with a cut-off period of 100 minutes has a very different meaning in situations where $\Delta t = 1$min or $\Delta t = 10$min. In the latter case, the effective cut-off period $\frac{100\text{min}}{10\text{min}} = 10\Delta t$ is dangerously low for detrending (see Figure S6 right panels), whereas for the much faster sampling a cut-off period of $\frac{100\text{min}}{1\text{min}} = 100\Delta t$ is fine for most if not all cases.

White noise has most of its energy concentrated in the high frequencies such that smoothing, even with the small cut-off period of four sampling intervals $T_c = 4\Delta t$, already removes half of all energy and gives 100% gain to the remaining periods, i.e. $\gamma_S(4\Delta t) = 2$. On the contrary, removing low-frequency components via detrending has very little effect, e.g. even keeping only the periods which are smaller than $T = 10\Delta t$ only gives a 25% gain, i.e. $\gamma_D(10\Delta t) = 1.25$. At higher cut-off periods, smoothing can lead to almost astronomical gains. For example, a $\gamma_S(20\Delta t) \times 100 = 1000\%$ (10-fold) increase in the average Wavelet power is obtained for $T_c = 20\Delta t$.

Compared to white noise AR1 noise has less energy located in the high frequency region of the spectrum. Thus, the (area-)ratio of $S$ and $D$ is different and ultimately the gains from smoothing get closer to the gains attained from detrending. For not too high values of $\alpha$, smoothing will practically always introduce more spurious oscillations than detrending. However, care has to be taken if the signal is only poorly sampled. In case the expected period is only a few multiples of the sampling interval, e.g. $T_{signal} \approx 7\Delta t$, the practitioner might be tempted to choose a very small cut-off period, e.g. $T_c = 10\Delta t$. As can be seen in Figure S6, this can easily introduce a substantial gain and hence spurious oscillations.

Pink noise stands out once more, as both detrending and smoothing can in principle equally introduce spurious oscillations, depending strongly on the value of $\beta$. In the analysis of signals where pink noise could be a substantial component, special care has to be taken. Decreasing the sampling interval and choosing very large cut-off periods for detrending is advisable.

#### A.4. Amplitude estimation

The normalization to unit energy, as defined in section A.1, allows together with the variance normalization of the signal for the extremely useful statistical interpretation of the Wavelet power. However, the instantaneous amplitude is a quantity of potential interest, especially for the practitioner. Therefore in other implementations of the continuous Wavelet transform, see e.g. Leise [2013], the Wavelets are defined so to yield the *amplitude spectrum*. A possible definition is, to set the maximum amplitude of the Wavelet in the frequency domain at the center frequency to $\widehat{\Psi}_{s,\tau}(\omega_c) = 2$. This ensures that for $f(t) = A_0\ cos(\omega t)$, the transform fulfills $|\mathcal{W}[f](t, \omega_c/\omega)| = A_0$ (see Lilly and Olhede [2010]).

As the Wavelet transform itself is a linear operator, and $|az|^2 = a^2|z|^2$ for $a \in \mathbb{R}$ and $z \in \mathbb{C}$, a simple re-scaling of the Wavelet power spectrum yields the amplitude spectrum. For the Morlet Wavelet the scaling factor is given by

$$\kappa(s) = \sqrt{\frac{2}{s}}\sigma\pi^{-1/4} \tag{31}$$

such that every scale (or period) has its own scaling factor.

### B. Supplementary Figures

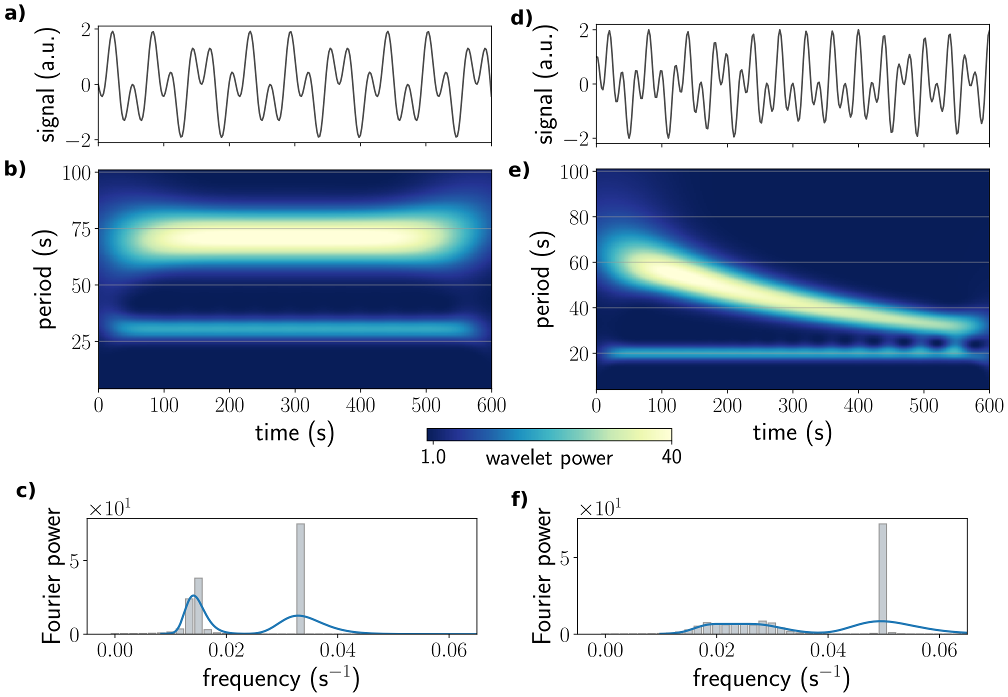


Figure S1: **Harmonic decomposition with Wavelets** a) Synthetic signal composed of two harmonic components with $T_1$ = 40s and $T_2$ = 90s. b) Wavelet power spectrum shows two bands with constant period. c) The corresponding Fourier power spectrum (bars), and the time averaged Wavelet power spectrum (solid line). d) The chirp signal from the main text figure 1a, augmented by a fast harmonic component with period $T_{fast} = 20s$ ($f_{fast} = 0.05Hz$) This linear superposition leads to a complicated waveform in the time domain. e) The Wavelet power spectrum clearly resolves the sweeping through periods from $T_2$ = 70s ($f_2 \approx$ 0.014Hz) to $T_1$ = 30s ($f_1 \approx$ 0.033Hz), and additionally also captures the fast harmonic component with constant period $T_{fast} = 20s$. f) The Fourier power spectrum can't resolve the sweeping instantaneous periods, but shows a prominent peak at $f_{fast} = 0.05Hz$.

Ravn, M. O. and Uhlig, H. (2002). On adjusting the hodrick-prescott filter for the frequency of observations. Review of economics and statistics, 84(2):371–376.

Torrence, C. and Compo, G. P. (1998). A practical guide to wavelet analysis. Bulletin of the American Meteorological society, 79(1):61–78.

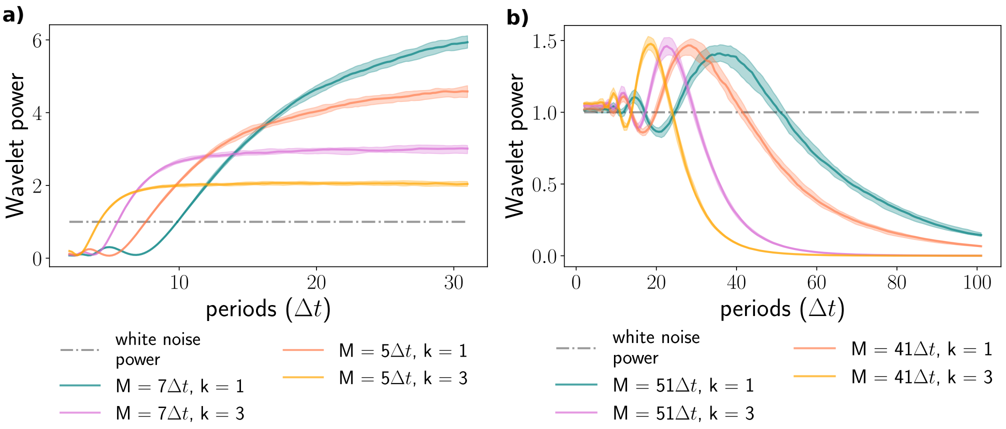


Figure S2: **Filter response of the Savitzky-Golay (or LOESS) filter** Window sizes (M) and polynomial order (k) as indicated. Numerical computation by filtering and Wavelet transform an ensemble of 50 long (25.000 sample points each) white noise trajectories. Higher polynomial order moves the effective cut-off periods towards smaller periods and decreases the roll-off of the filter. a) Smoothing (low-pass) response. b) Detrending (high-pass) response.

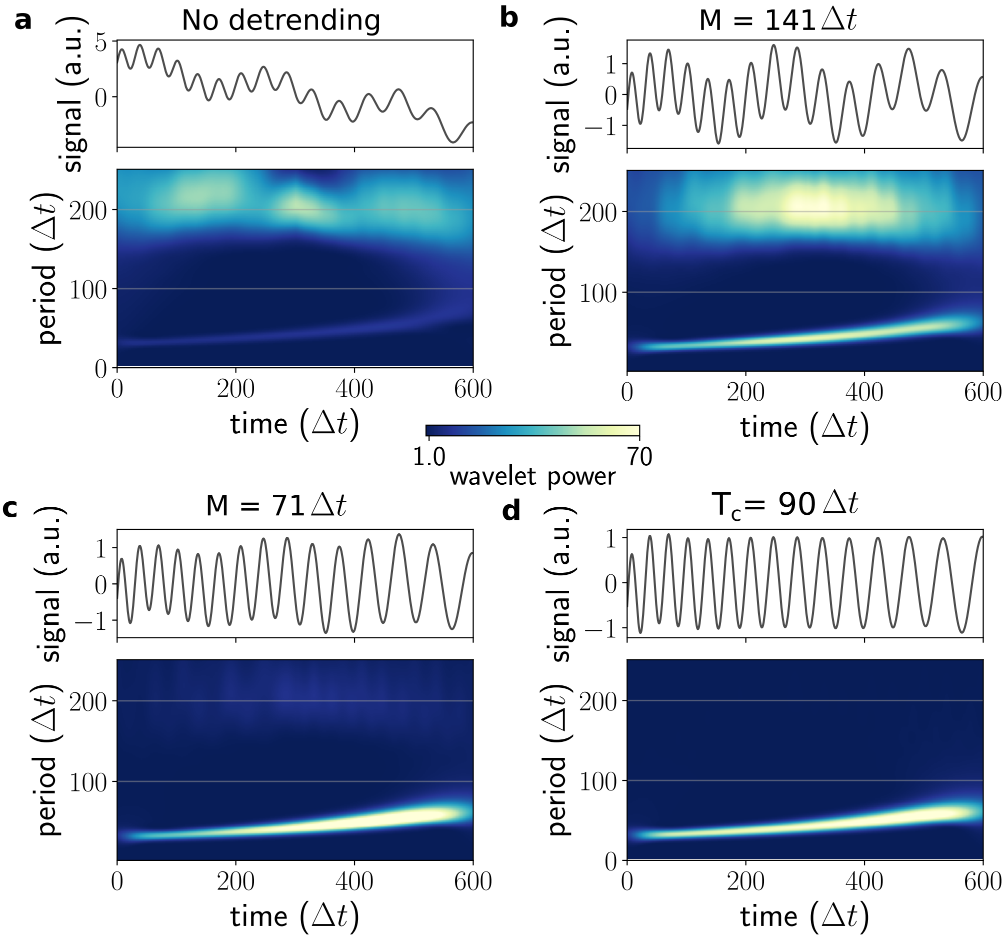


Figure S3: **Wavelet spectra for different detrending** Synthetic signal from Figure3 of the main text. a) Without detrending, the actual signal carries only very little power. b) Detrending with a very wide moving average filter removes the linear trend, however the slow oscillatory trend component still dominates the signal. c) The optimal moving average filter almost completely removes the trend. Slight modulations are still visible in the filtered signal. d) The sinc filter completely removes the trend.

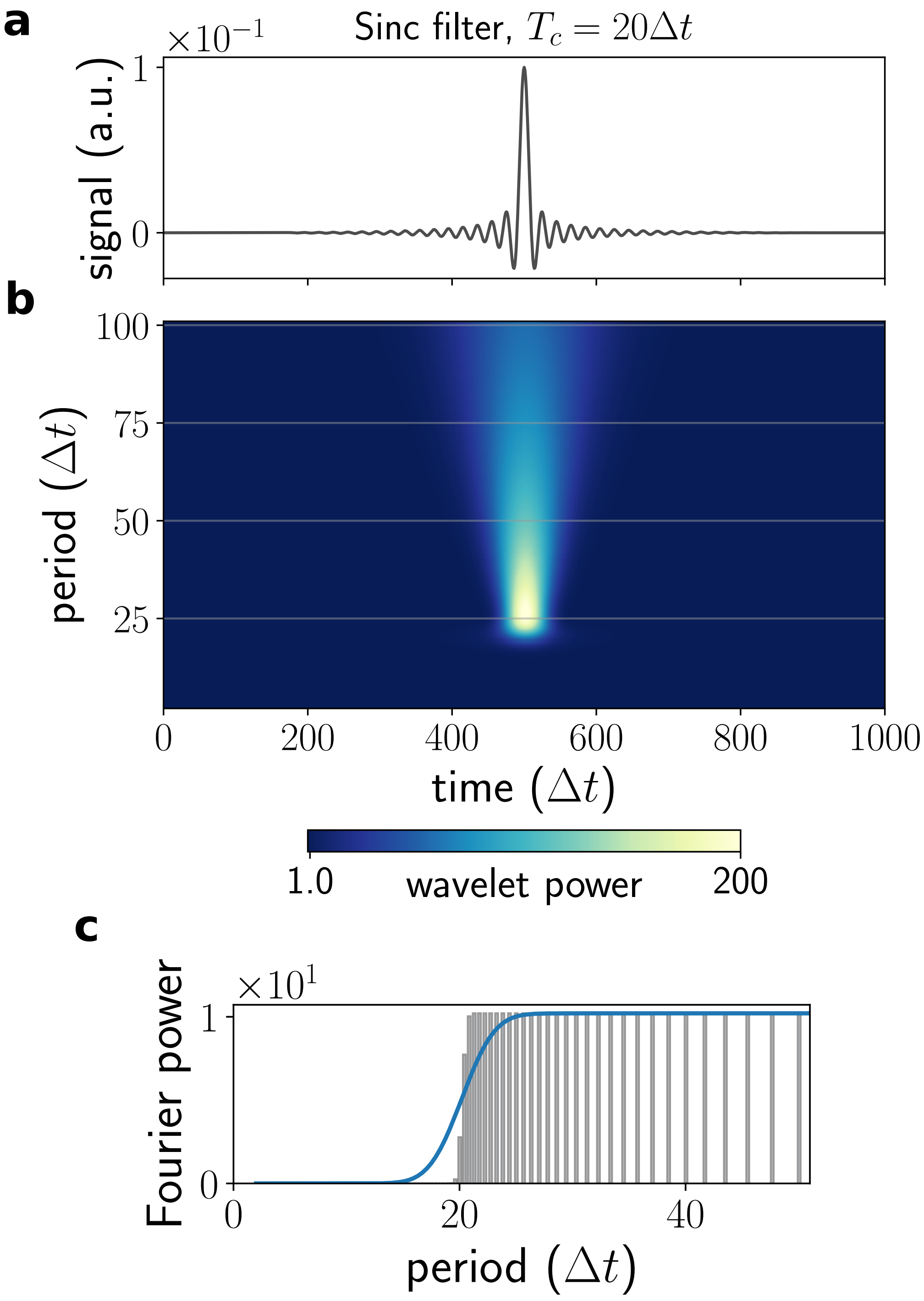


Figure S4: **The sinc filter** a) Time representation of the sinc filter. b) Wavelet power spectrum. c) Fourier power spectrum (bars) and averaged Wavelet power spectrum (solid line).

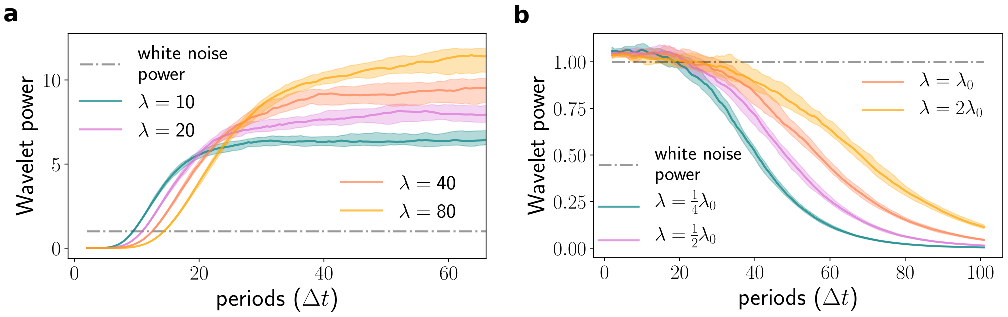


Figure S5: **Filter response of the Hodrick-Prescott filter** Different values of $\lambda$ as indicated. Numerical computation by applying the filter and Wavelet transform an ensemble of 50 long (25 000 sample points each) white noise trajectories. a) Smoothing (low-pass) response. b) Detrending (high-pass) response. The reference value $\lambda_0 = 0.05 * \left(\frac{T}{\Delta t}\right)^4$ for $T = 24\Delta t$ is taken from the literature (Ravn and Uhlig [2002]).

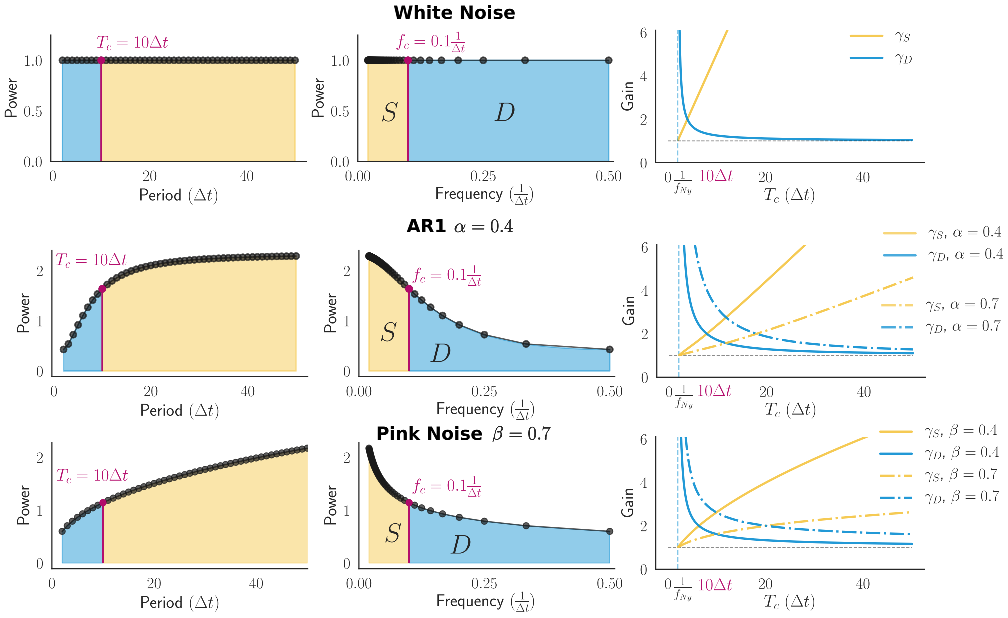


Figure S6: **Gains introduced by smoothing and detrending with a sinc-filter for three different types of noise** See also supplementary section A.3. *Left panel*: Period representation of the power spectra. An example cut-off period $T_c = 10\Delta t$ for the sinc filter is marked. Smoothing (detrending) retains the high (low) periods in the spectrum. *Middle panel*: Frequency representation of the power spectra. The parts of the spectrum which remain after smoothing (detrending) are indicated as $S$ ($D$). The Fourier bins given by the sampling interval: $\frac{1}{i\Delta t}$, with $i = 2, ..., N$ are indicated as black dots. The ratio of $S$ to $D$ changes significantly for the presented spectra. In the period representation this ratio has no physical meaning. *Right Panel*: The gain functions for smoothing ($\gamma_S$) and detrending ($\gamma_D$) as a function of the cut-off period ($T_c$). Both AR1 ($\alpha$) and pink noise ($\beta$) have a free parameter. Two curves are plotted each for smoothing and detrending. The smallest available cut-off period is given by $1/f_{Ny}$, with $f_{Ny}$ the denoting the Nyquist frequency.

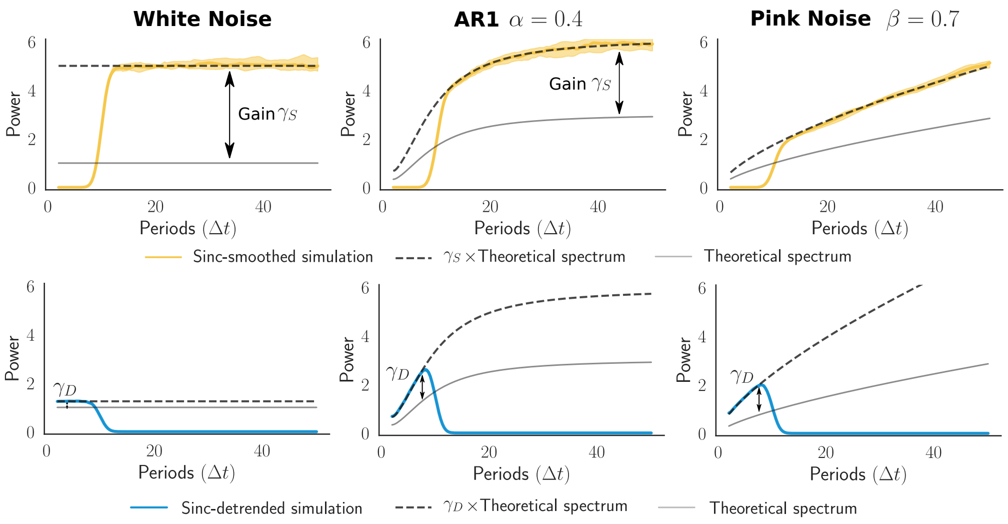


Figure S7: **Comparison of the predicted filtered power spectra with simulations** As outlined in section A.3, the theoretical power spectra are scaled with the analytically calculated gains for the sinc cut-off period of $T_c = 10\Delta t$. An ensemble of 50 realizations for each of the noise processes was sinc-filtered ($\gamma_S$ - smoothing, $\gamma_D$ - detrending), and their median and quartiles of the time-average Wavelet spectra plotted against the predictions. The ratio of the gains changes considerably depending on the type of noise, see also FigureS6.

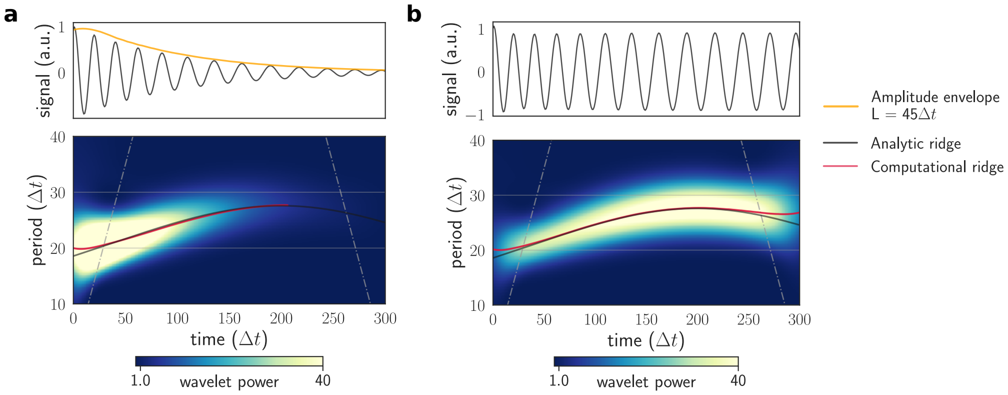


Figure S8: **Amplitude decays lead to impractical power values** a) A noise free signal with an exponential decaying amplitude envelope. The spectrum shows a strong decline in power, leading to an incomplete computational ridge. b) Amplitude normalized signal and its Wavelet spectrum. After removal of the envelope, the signal can be optimally traced with a ridge in the spectrum.

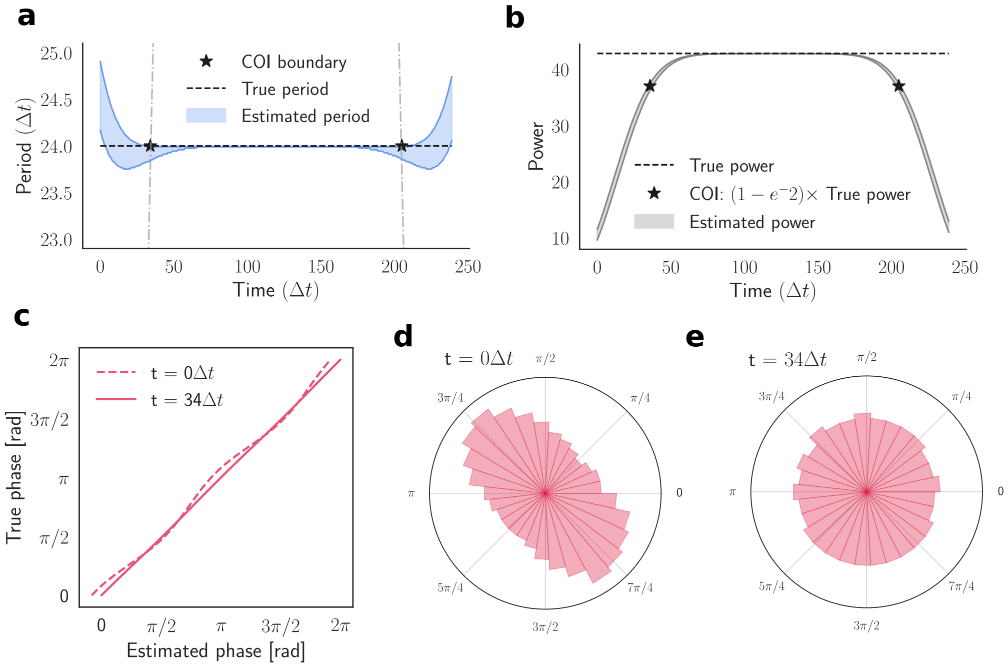


Figure S9: **Edge effects and the cone of influence (COI)** Wavelet analysis of an ensemble of phase shifted cosines: $f(t, \phi) = cos(\omega t + \phi)$, with $\omega = 2\pi/24\Delta t$ and $\phi = [0, 2\pi)$. a) Quartiles of estimated periods, inside the COI the maximal deviation from the true period is $\approx 4\%$. The width of the COI is given by the e-folding time of the Morlet Wavelet at the respective period: $COI_{width}(T) \approx 1.43T$. Peaks in the Wavelet spectra at the respective periods should be wider than this decorrelation time. Otherwise it's just a single spike in the data. b) Quartiles of estimated powers, the power drops significantly inside the COI. At the boundary of the COI, the deviation between true and estimated power is given by the factor $e^{-}2$, see also Torrence and Compo [1998]. c) Phase map of estimated vs. true phases for the initial time point ($t = 0\Delta t$) and at the boundary of the COI $t = 34\Delta t$. d) Bimodal phase distribution at $t = 0\Delta t$. This bimodality is induced solely by the edge effects inside the COI. The true distribution is uniform. e) Recovery of the uniform phase distribution at $t = 34\Delta t$.

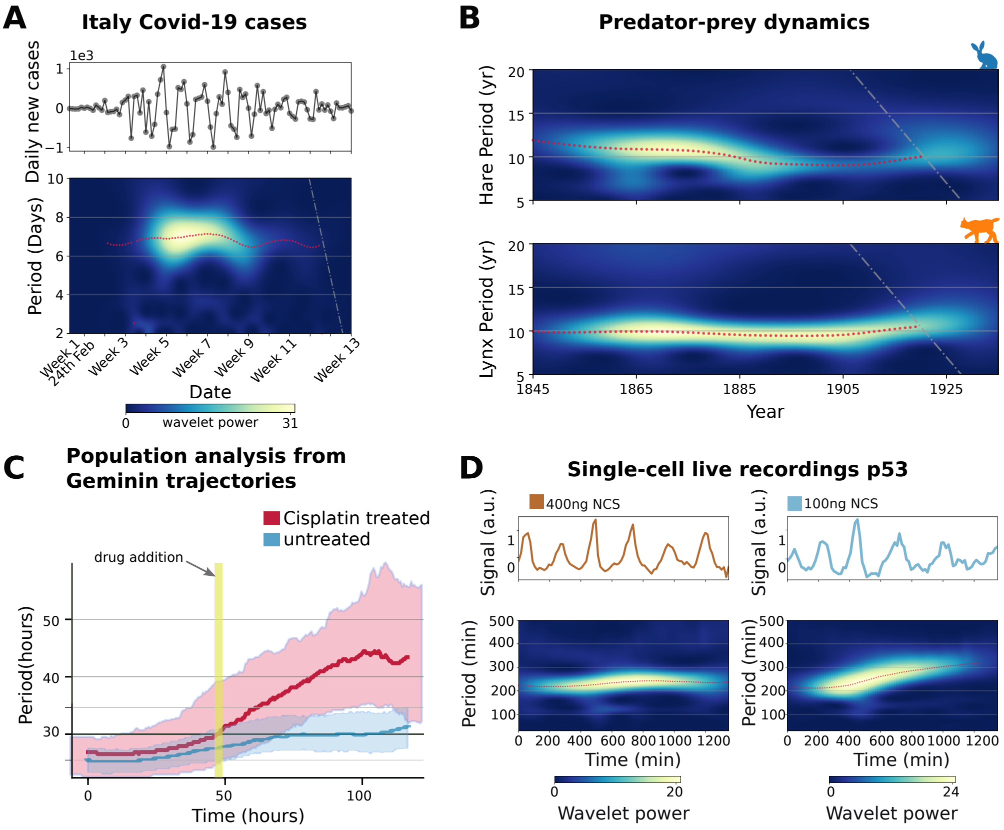


Figure S10: `pyBOAT` **applications** Additional and intermediate results for the analysis of the example datasets shown in the main text Figure 5. A) Detrended signal and wavelet spectrum for the Covid-19 data from Italy. B) Wavelet power spectra for the hare and lynx signals. A 10 year rhythm is clearly discernible for both species. C) Time-dependent period distribution of the U2OS cells, with the experimental conditions as indicated. The cisplatin treated cells slow down their cell cycle significantly on the populational level. D) Two representative single-cell p53 signals, highly stimulated (left) and medium stimulated (right).

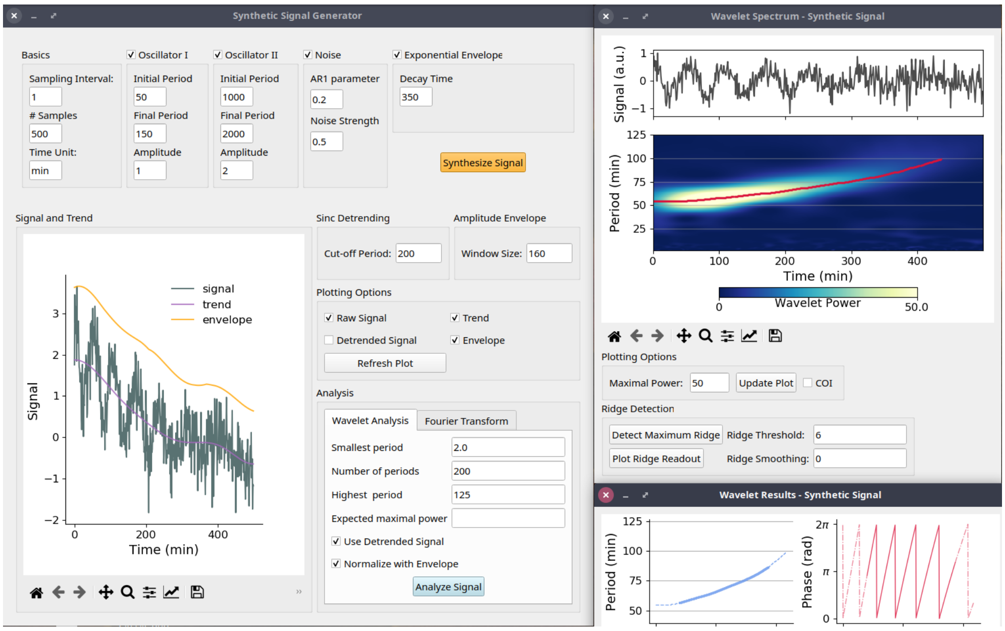


Figure S11: **pyBOAT synthetic signal generator** Screen capture of the synthetic signal generator. The main oscillatory component in this example here is a chirp with slowing down period (50min - 150min). To simulate non-linear trends, a second oscillator ('Oscillator 2') can be activated and set to very long periods. AR1 noise and an amplitude envelope can also be added to simulate challenging signal components.
